# A Coronaviral Pore-Replicase Complex Drives RNA Synthesis in Double Membrane Vesicles

**DOI:** 10.1101/2024.06.18.599549

**Authors:** Anan Chen, Ana-Mihaela Lupan, Rui Tong Quek, Stefan G. Stanciu, Mihaela Asaftei, George A. Stanciu, Kierra S. Hardy, Taciani de Almeida Magalhães, Pamela A. Silver, Timothy J. Mitchison, Adrian Salic

## Abstract

Coronavirus-infected cells contain double-membrane vesicles (DMVs) that are key for viral RNA replication and transcription, perforated by hexameric pores connecting the vesicular lumen to the cytoplasm. How pores form and traverse two membranes, and how DMVs organize RNA synthesis, is unknown. Using structure prediction and functional assays, we show that the non-structural viral membrane protein nsp4 is the key DMV pore organizer, spanning the double membrane and forming most of the pore lining. Nsp4 interacts with nsp3 on the cytoplasmic side and with the viral replicase inside the DMV. Newly synthesized mRNAs exit the DMV into the cytoplasm, passing through a narrow ring of conserved nsp4 residues. Steric constraints imposed by the ring predict that modified nucleobases block mRNA transit, with broad spectrum anti-coronaviral activity.

## Main Text

Coronaviruses have large single stranded (ss) RNA genomes (up to 33 kb) [1] that encode a polyprotein followed by four structural and several accessory proteins. The polyprotein is cleaved into sixteen non-structural proteins (nsps) [2] that play key roles in the viral takeover of the infected cell. Nsps 7,8,9,12 and 13 form the RNA-dependent RNA polymerase complex (RdRp or replicase), which first copies and then transcribes the viral genome, of which Nsp12 in the catalytic subunit [3–5]. The three transmembrane nsps 3,4 and 6 remodel endoplasmic reticulum (ER) membranes into characteristic double-membrane vesicles (DMVs). DMVs are thought to play a central role in viral RNA synthesis and to protect the dsRNA replication intermediate from recognition by cytoplasmic innate immune sensors [6–9]. DMVs have an inner lumen containing filament-like structures [10], identified as double stranded RNAs (dsRNAs) that represent viral genome replication intermediates [10, 11]. The lumen is connected to the host cytoplasm via six-fold symmetric pores that contain nsp3 [12, 13] and are thought to mediate the exchange of molecules between the two compartments. Other viral nsps, such as nsp4 and nsp8, have been localized in the proximity of DMVs or in adjacent membranous structures, by light microscopy or immuno-electron microscopy [11, 14, 15]. However, the molecular nature of the DMV pores, the relationship between the replicase and DMVs, and the precise role of DMVs in viral RNA synthesis remain unknown.

Pioneering cryo-EM tomography studies in mouse hepatitis virus (MHV)-infected cells showed that six copies of nsp3 form the cytoplasm-facing aspect of the DMV pore [12]. However, nsp3 contains only two transmembrane helices (TMs), suggesting that a nsp3 hexamer alone would be insufficient on its own to delineate a pore that is more than 2 nm in diameter at its narrowest and traverses two phospholipid bilayers that are 10-11 nm apart [12]. We reasoned that nsp4, which has six TMs, is a good candidate for the main coronaviral pore-forming protein. ColabFold multimer [16, 17] predicted with high confidence that nsp4 assembles into a symmetric hexameric complex with a central tunnel (fig S1A). This predicted structure is highly conserved among coronavirus species, despite modest sequence conservation (fig S1B). The tunnel’s smallest diameter is 1.9nm (Fig 1E), slightly less than the diameter measured for DMV pores in cells [12].

**Fig. 1.**
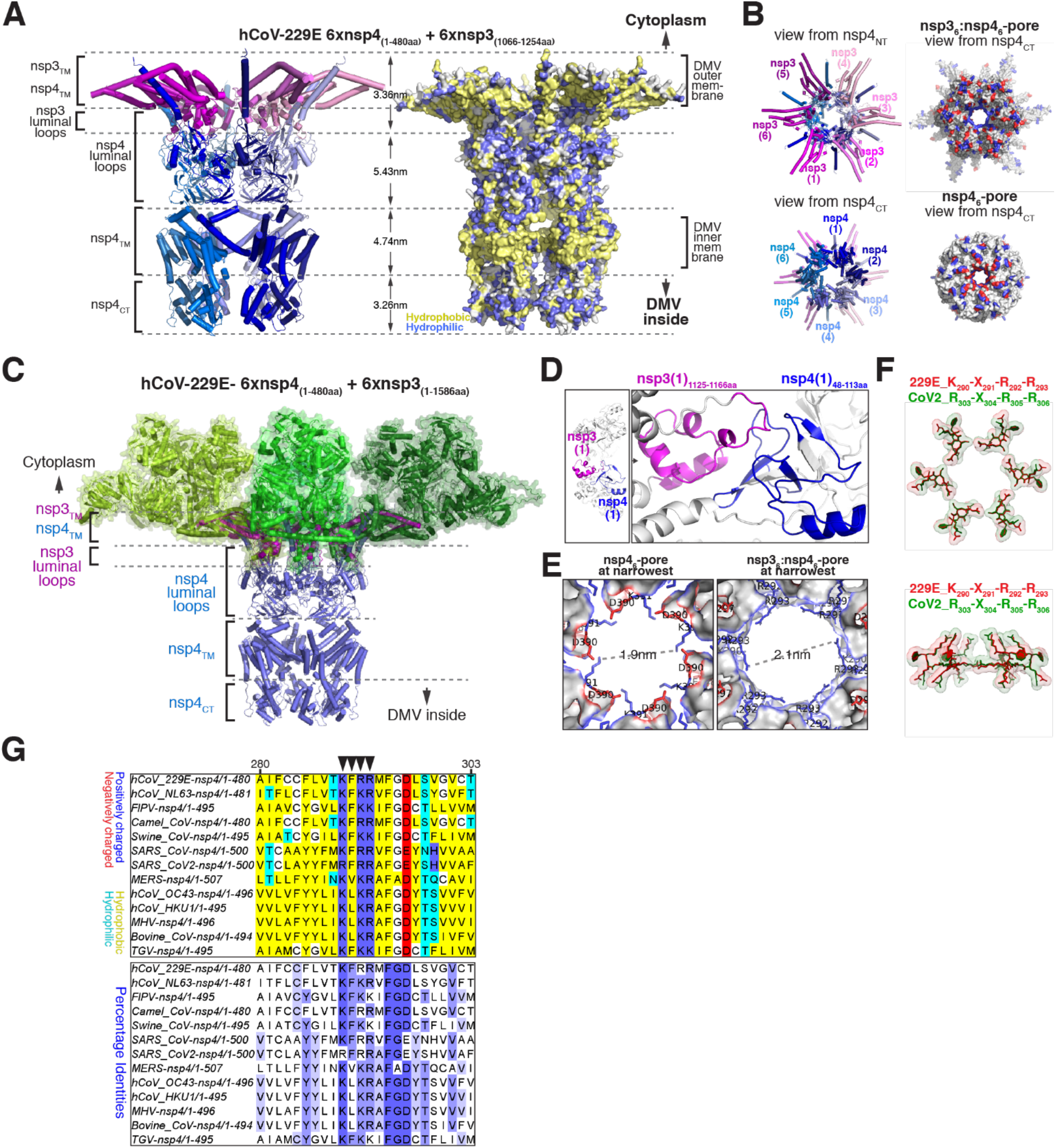
A coronaviral nsp3-nsp4 complex defines the DMV molecular pore. **(A)** Predicted structure of a six-fold symmetric complex between hCoV-229E nsp3 (residues 1066-1254) and full-length nsp4, seen from the side. TM = transmembrane domains; NT = N-terminus; CT = C-terminus. Left: ribbon diagram; right: molecular surface showing hydrophobic (yellow) and hydrophilic (blue) residues. Distances between domains are indicated. **(B)** As in (A), but showing top and bottom views of the complex, respectively. Left: ribbon diagram; right: molecular surface showing positive (blue) and negative (red) residues. **(C)** As in (A), but including the predicted structure of full-length nsp3 (green). **(D)** Close-up view of the interaction between the inter-membrane domains of hCoV-229E nsp3 and nsp4. **(E)** Predicted narrowest portion of the pore formed by hCoV-229E nsp4 alone (left) or in complex with nsp3 (right). The diameter of the constriction and the conserved residues delineating it are indicated. **(F)** As in (E), but showing the overlay of the conserved pore constriction residues in hCoV-229E (red) and SARS-CoV2 (green). **(G)** Sequence alignment of nsp4 homologs from various alpha or beta-coronaviruses, showing the conserved K-X-R/K-R/K pore motif (arrows).

Since nsp4 interacts with nsp3 [18–20], we next obtained a high-confidence model of a nsp3-nsp4 hexamer, using a C-terminal fragment of nsp3 that is sufficient for DMV formation, which includes the two TMs and the intervening luminal domain (Fig 1A, 1B). We then aligned six copies of the predicted full-length nsp3 (Fig 1C; fig S1C-E), which fit into the model without steric clashes. The resulting six-fold symmetric nsp3-nsp4 complex (Fig 1C) is strikingly similar in size and shape to the electron density of the DMV pore determined by cryo-EM tomography [12]. The predicted pore has a continuous central tunnel, the narrowest portion of which is now increased to 2.1-2.3 nm (Fig 1B, 1E, 1F; fig S1F), in excellent accord with measurements in virus-infected cells [12]. This narrowest portion is delineated by a ring of eighteen positively charged residues contributed by the six nsp4 molecules (Fig 1E, 1F; fig S1F); these residues form a consensus (-K/R_290_-X_291_-R/X_292_-R/K_293_-) that is highly conserved among nsp4 homologs (Fig 1F, 1G).

A major unanswered question is how the nsp3/4 pore spans the two concentric, closely-spaced membranes. In our model, each nsp4 monomer uses the last five of its six TMs to traverse the inner membrane (Fig 1A, 1C). The ER-luminal domain of nsp4 forms most of the portion of the pore spanning the intermembrane space (Fig 1A). The model further predicts a high-confidence interface between the ER-luminal domains of nsp3 and nsp4 (Fig 1D), consistent with longstanding binding data [18–20]. The remaining single N-terminal TM of nsp4 lies on the opposite side of the molecule from the other five TMs, both when nsp4 is modeled alone (fig S1A) or in complex with nsp3 (Fig 1A), suggesting that nsp4 plays a key role in spanning the double membrane. This arrangement is in sharp contrast to a recent prediction that has nsp4 residing entirely in the inner DMV membrane [13]; furthermore, our model is the first to account for both nsp3-nsp4 co-assembly and the crossing of both DMV membranes. The N-terminal TM of each nsp4 monomer, together with the two TMs of each nsp3 molecule, form the portion of the pore that spans the outer membrane, while the soluble N-and C-terminal domains (NTD and CTD) of nsp3 form the entire cytoplasm-facing portion of the pore (Fig 1C; fig S1D). A key prediction of our model is that both N and C-termini of nsp3 face the cytoplasm while those of nsp4 face the cytoplasm and the inner lumen of the DMV, respectively (Fig 1C; fig S1D).

To test the predicted topology of the nsp3-nsp4 complex, we first used a reconstituted cellular system in which co-expression of nsp4 with truncated nsp3 generates abundant synthetic DMVs (∼100nm in diameter) in the absence of viral infection (fig S2A). Nsp3 and 4 show strong colocalization and interaction (fig S2A, S2B) in this minimal system. We then used differential permeabilization and immunofluorescence to probe antibody accessibility of tags attached to the N and C-termini of nsp3 and nsp4. In the absence of nsp3, both N and C-terminal tags on nsp4 were equally accessible to antibodies (Fig 2B), whether the cells are treated with low amounts of digitonin, which permeabilizes the plasma membrane but not intracellular membranes, or with Triton-X100 (TX-100), which permeabilizes all membranes (Fig 2B, 2H; fig S3A). As expected, a nuclear-localized protein was accessible to antibodies only after TX-100, but not digitonin, permeabilization (fig S3B, S3B). Thus, both nsp4 termini face the cytoplasm when the protein is expressed alone. However, when synthetic DMVs were induced by co-expressing nsp3, the nsp4 C-terminus became inaccessible with digitonin, while accessibility of the N-terminus was unchanged (Fig 2C, 2D, 2G; fig S3F, S3G). Importantly, the N-and C-termini of nps3 were both accessible with digitonin permeabilization (Fig 2C, 2D, 2G; fig S3H, S3I). Similar results were obtained using synthetic DMVs immunopurified via a C-terminal tag on nsp3 (Fig E2C-F; fig S3D, S3E). Next, we tested antibody accessibility in viral DMVs formed during hCoV-229E infection, using affinity-purified antibodies against the NTD of nsp3 and the CTD of nsp4, and obtained similar results to those in synthetic DMVs (Fig 2h-k). Together, these results confirm the predicted topology of the nsp3-nsp4 molecular pore complex: both termini of nsp3 face the cytoplasm, while the N-and C-termini of nsp4 face the cytoplasm and the inner DMV lumen, respectively. The fact that nsp4 spans the two membranes, while both its termini face the cytoplasm when expressed alone, suggests that nsp4 plays a central role in pore assembly and DMV closure.

**Fig. 2.**
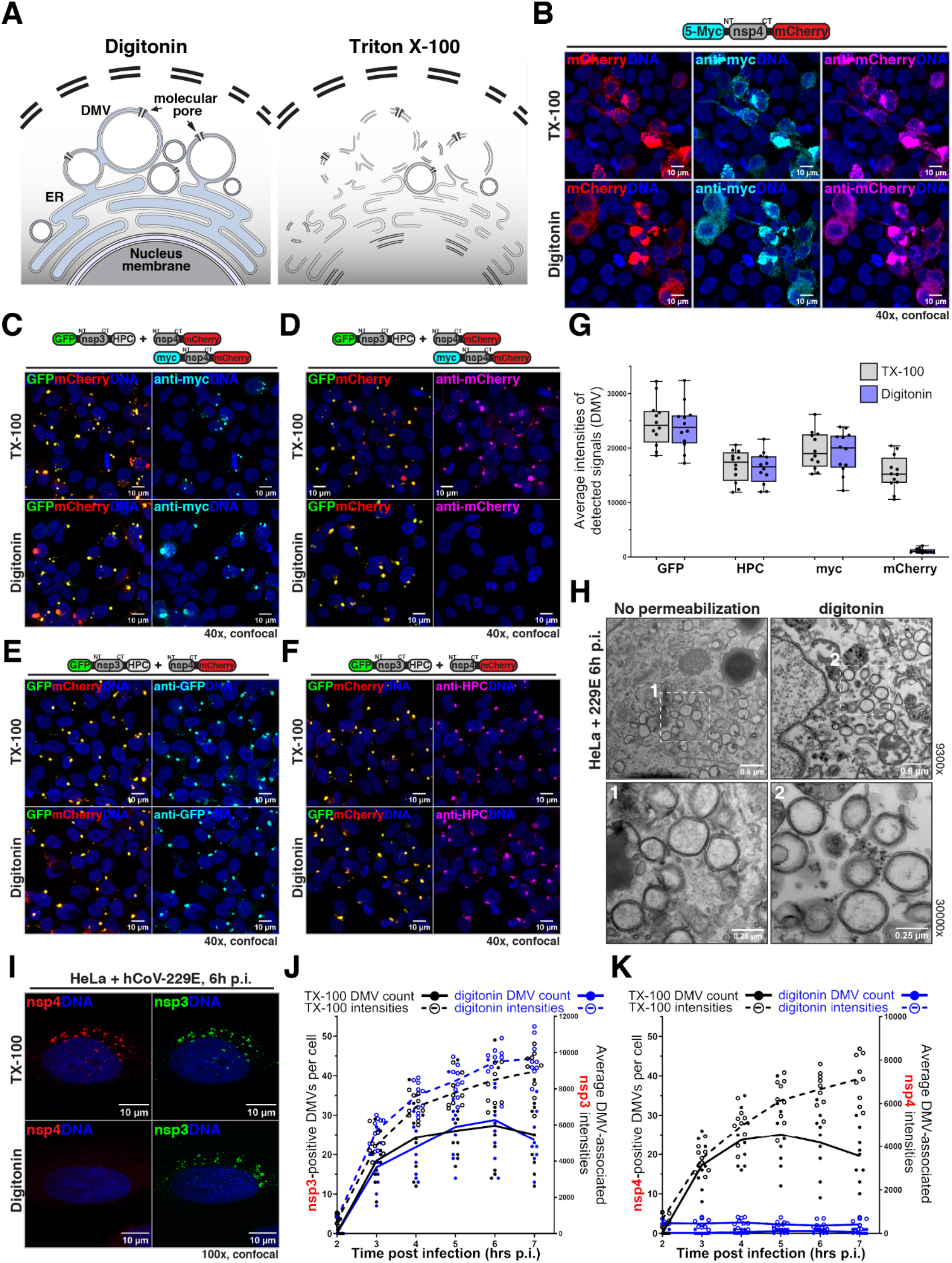
Nsp4 spans both DMV membranes, extending from the cytoplasm to the lumen. **(A)** Differential antibody accessibility assay. Fixed cells permeabilized with digitonin or Triton-X100 are subjected to immunofluorescence staining. **(B)** In cells expressing myc-nsp4-mCherry, both myc and mCherry tags face the cytoplasm. **(C, D)** As in (B), but with cells co-expressing eGFP-nsp3ΔN-HPC and myc-nsp4-mCherry. The myc tag now faces the cytoplasm, while the mCherry tag faces the DMV lumen. **(E,F)** As in (C), but stained with indicated antibodies. **(G)** Quantification of experiments in (C-F). Four non-overlapping fields were analyzed per condition and each condition was repeated thrice. (*N*=3, n=12). **(H)** Transmitted electron microscopy imaging of hCoV-229E-infected cells showing viral DMVs at 6 hours post infection. Close-ups of the outlined regions are shown at the bottom. **(I)** As in (H), but cells were differentially permeabilized and co-stained with antibodies against nsp3-NT and nsp4-CT. Nsp3-NT faces the cytoplasm while nsp4-CT faces the DMV lumen. **(J, K)** As in (I), showing quantification of nsp3 (J) or nsp4 (I) differential accessibility at different time points post infection. Three non-overlapping fields of view were analyzed per condition and each condition was repeated thrice. (*N*=3, n=9).

We extended the accessibility analysis to localize dsRNA in cells infected with hCoV-229E using affinity-purified antibodies against viral proteins and a novel probe based on protein kinase R (PKR) that detects dsRNA with high sensitivity (Fig 3; fig S4). dsRNA and viral proteins were detected as early as 2 hours after infection (Fig 3A, 3B; fig S4B-F). At all time after infection, dsRNA and the core RdRp subunit nsp7 were inaccessible to protein probes under digitonin permeabilization conditions (Fig 3A, 3B; fig S4B, S4D), similar to the C-terminus of nsp4, while two other RdRp subunits, nsp8 and nsp9, were weakly accessible (Fig 3A; fig S4A, S4E, S4F); all these components were accessible following TX-100 permeabilization, showing that the viral dsRNA and the replicase are inside DMVs. Since dsRNA is the first viral RNA species synthesized upon infection [21][22] the replicase must be rapidly enclosed inside DMVs following polyprotein translation. In contrast, newly synthesized DMV-associated RNA, metabolically labeled with 5’-ethynyl-uridine (EU) [23, 24] and detected by click reaction with biotin, was partially accessible to fluorescent streptavidin (Fig 3B; fig S4C). Similarly, enzymatic digestion of EU-labeled viral RNA was partial in the presence of digitonin and complete with TX-100 (fig E5A-D). Based on sensitivity to RNase A, which digests ssRNA but not dsRNA (fig E5A, E5B), the newly synthesized DMV-associated RNA is single stranded. We also examined viral RNA synthesis using a run-off assay in digitonin-permeabilized cells (fig E5E). Under these conditions, nascent DMV-associated RNA is protected from nuclease digestion (fig E5F, E5G). Together, these data indicate that, while the dsRNA intermediate and the replicase complex are inside DMVs, viral transcripts spread from inside DMVs to the cytoplasm.

**Fig. 3.**
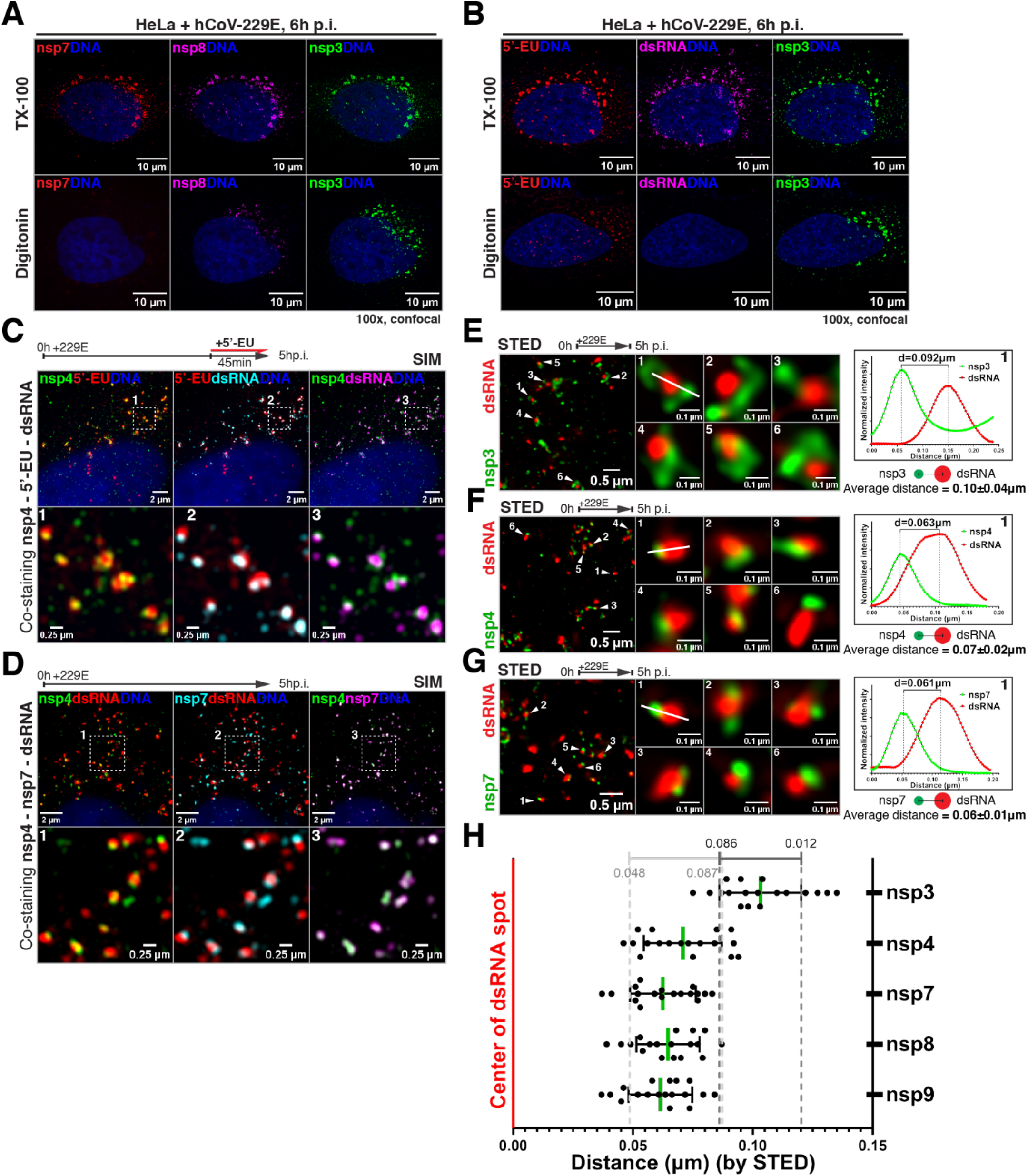
The viral replicase localizes inside DMVs, in close proximity to nsp4. **(A and B)** hCoV-229E-infected cells labeled with EU were fixed and permeabilized, then (A) stained with fluorescently-labeled antibodies against nsp7, nsp8 and nsp3, or (B) reacted by click chemistry with biotin-azide, followed by co-staining with fluorescent streptavidin, dsRNA probe, and anti-nsp3 antibodies. Nsp7 and dsRNA are inside DMVs, while EU-labeled transcripts are both inside DMVs and cytoplasmic. **(C)** As in (A, B), but with staining for nsp4 and imaging by SIM. **(D)** As in (C), but with co-staining for nsp4, nsp7 and dsRNA. **(E-G)** hCoV-229E-infected cells were fixed, permeabilized and co-stained for (E) nsp3 and dsRNA, (F) nsp4 and dsRNA, or (G) nsp7 and dsRNA, followed by imaging by STED microscopy. Middle panels show zoomed-in views of DMVs on the left. The white line in middle panel 1 was used for linescan analysis (right panel), to measure distance between nsps and dsRNA. **(H)** Quantification of center-to-center distance between nsps and dsRNA in (G). Eighteen zoomed-in images from 3 non-overlapping fields were analyzed for each group (n=18). Nsp4, nsp7, nsp8 and nsp9 localize closer to dsRNA, while nsp3 is further away.

We next used structural illumination microscopy (SIM) and stimulated emission depletion microscopy (STED) to pinpoint the location of key components involved in viral RNA synthesis. We focused on single DMVs (dsRNA signal diameter ≤ 250nm) undergoing active transcription early in the infection (fig E6A-D), as identified by EU incorporation (Fig 3C). Components of the pore (nsp3 and nsp4) and replicase (nsp7, nsp8 and nsp9), as well as the dsRNA localized within a larger and more diffuse EU-labeled RNA signal (Fig 3C; fig S6I, S7A-C). Nsp4 shows a high degree of co-localization with nsp7 and nsp8 throughout viral infection, while nsp3 colocalizes to a lesser degree (Fig 3D; fig S6I, S7D, S7E). Nsp4, nsp7, nsp8 and nsp9 always lay closer to the dsRNA signal, while nsp3 was further away (Fig 3E-H; fig S6E, S6F); this is consistent with the predicted localization of the N-terminus of nsp3 and the C-terminus of nsp4 to the DMV surface and inner lumen, respectively (Fig 1G). These data are also consistent with results obtained by horseradish peroxidase (HRP)-catalyzed proximity labeling, which place nsp4, nsp7, nsp8 and dsRNA close together (fig S4G-K). The degree of colocalization between nsps, and between nsps and dsRNA, decreases as infection progresses (fig E6G, E6H); the reason for this is currently unknown and might involve overlap of signals at the light level with increased density. This might explain why past studies did not observe a close spatial relationship between replicase and dsRNA in DMVs [11]. Together, our high-resolution fluorescence imaging suggests that the DMV pore and the replicase form a complex.

To test this hypothesis, a DMV-enriched fraction from infected cells (Fig 4A, 4B) was detergent-solubilized and subjected to immunoprecipitation and Western blotting. As shown in Fig. 4C, nsp4, nsp7 and nsp8 are precipitated with anti-nsp3 antibodies, while nsp3, nsp4 and nsp7 are precipitated with anti-nsp8 antibodies. Furthermore, mass spectrometric analysis shows that nsp9, nsp12 and nsp13 are also precipitated with anti-nsp3 and anti-nsp8 antibodies (fig S8A, S8B). These results indicate that a pore-replicase complex exists inside DMVs. Since nsp4 is the lumen-facing pore subunit, it is likely that it is involved in contacting the replicase. We find that nsp4 interacts with nsp8 alone or with a nsp7-nsp8 complex (fig S8E), corroborating the prediction of a nsp4-nsp8 hexamer by ColabFold (fig S8F); however, whether the replicase is connected to the pore through nsp8 alone, of by a more extended molecular interface, possibly also involving RNA molecules, remains to be determined.

**Fig. 4.**
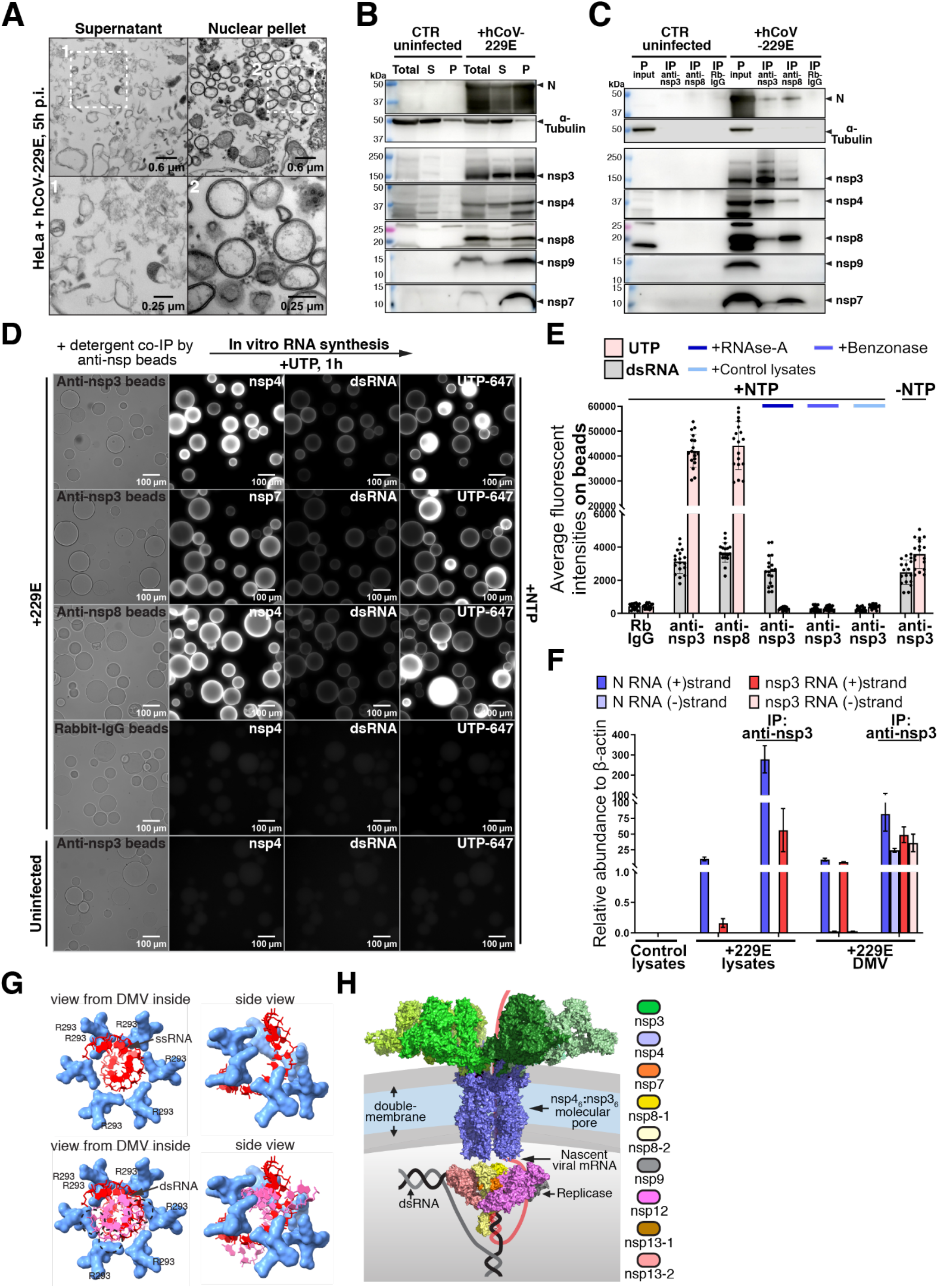
Viral RNA synthesis by a DMV replicase-pore complex. **(A and B)** hCoV-229E-infected cells were lysed, and nuclear pellets (P) or supernatants (S) were imaged by TEM (A), or analyzed by SDS-PAGE and immunoblotting (B); Total = total lysates; N = nucleocapsid. **(C)** As in (B), but solubilized (P) fraction was immunoprecipitated with anti-nsp3 or anti-nsp8 antibodies followed by immunoblotting. **(D)** As in (C), but immunoprecipitates on beads were incubated with NTPs and fluorescent UTP, then fixed and stained for dsRNA and nsps. **(E)** As in (D), but showing quantification of UTP and dsRNA intensities on beads. Six non-overlapping fields were analyzed per condition, and each condition was repeated thrice (*N*=3, n=18). **(F)** Positive (+) or negative (-) strand RNAs for N or nsp3 were measured by quantitative RT-PCR. RNA abundance was normalized to β-actin mRNA in each sample. Each condition was performed in triplicate (n=9). **(G)** Model of mRNA (top) and dsRNA (bottom) molecules translocating through the conserved pore constriction. The dsRNA molecule clashes sterically (dashed circles) with residues in nsp4. **(H)** Proposed model of the DMV pore-replicase complex transcribing a dsRNA template, with the nascent viral mRNA co-translocating through the pore to the cytoplasm.

Aside from the replicase, dsRNA and (-) strand viral RNAs also precipitate with anti-nsp3 antibodies (Fig 4D, 4F), and furthermore, dsRNA affinity isolated from the DMV-enriched fraction co-purifies with nsp3, nsp4, nsp8, nsp9, nsp12 and nsp13 (fig S8C). We thus tested whether the pore-replicase complex is active in RNA synthesis. When the complex immunopurified on anti-nsp3 beads is incubated with a ribonucleotide triphosphate (NTP) mix supplemented with fluorescent UTP, a strong fluorescent signal develops on the surface of the beads (Fig 4D, 4E; fig E8D). This signal is dependent on the presence of the NTP mix during incubation, and is sensitive to RNase A, indicating that it represents ssRNA transcribed on beads (Fig 4E; fig S8D). Interestingly, the amount of dsRNA remains constant on the beads during incubation (Fig 4D, 4E; fig S8D), consistent with the idea that it serves as template for the RdRp, similar to how a DNA duplex is template for RNA polymerase. Similar results were obtained when the pore-replicase complex was purified using anti-nsp8 beads (fig S8D). These results show that the pore-replicase complex is transcriptionally active, perhaps using dsRNA as template.

The conserved positively charged residues surrounding the narrowest portion of the nsp3/4 pore suggest an interaction with nascent viral mRNA as it exits the DMV lumen. Modeling predicts that the pore can accommodate one ssRNA molecule, but it is too narrow for dsRNA (Fig 4G). The steric constraint imposed by the Arg-rich ring in the pore predicts that bulky modifications of the nucleobases might interfere with mRNA passage through the ring, leading to an anti-viral effect. To test this, we pre-treated cells with N^6^-benzyl-adenosine and infected them with hCoV-229E. We observed a strong block of viral mRNA transcription, as assayed by EU incorporation in DMVs and viral assays, while adenosines with smaller N^6^ substituents had no effect (fig S5F, S5G, S9A, S9B, S9E-G). Importantly, all adenosine analogs compete with the incorporation of the “clickable” N^6^-propargyl-adenosine into total cellular RNA (fig S9C, S9D), indicating that the analogs were properly taken up by cells, converted to NTPs, and utilized in transcription. Inhibition of viral mRNA transcription by N^6^-benzyladenosine was comparable to that caused by the well-established RdRp inhibitor remdesivir [25, 26] (fig S9A, S9B). Interestingly, while remdesivir potently blocked both dsRNA and mRNA synthesis, N^6^-benzyladenosine had only a modest effect on dsRNA (fig S9A, S9B), consistent with an inhibitory effect on mRNA translocation through the pore instead of impeding the initial copying of the viral genome.

Our results suggest that coronaviral DMVs operate like simple dsRNA-based nuclei. Nsp4 plays a central role in the architecture of the pore and in bringing the two membranes together by spanning both of them. Nsp4 also recruits the replicase to the DMV lumen, most likely by an interaction between its C-terminal domain and the nsp8 subunit of the replicase. DMVs are probably inflated by synthesis of the dsRNA template trapped inside, as implied by the larger size of viral DMVs compared to synthetic ones [6][13]. Nascent viral mRNA is translocated co-transcriptionally through the pore to the cytoplasm, interacting with highly conserved Arg motifs in nsp4 that line the pore (fig S10A, S10B). It is unclear if other molecules pass through the pore besides exiting mRNAs; one possibility is that all necessary macromolecules are enclosed at the time of DMV formation. The NTPs required for RNA synthesis inside the DMV might freely enter through the pore or, alternatively, might pass through unidentified transporters. It appears that DMVs form rapidly after polyprotein translation and proteolytic processing, ensuring that dsRNA is immediately shielded from innate immune detection by cytoplasmic sensors. It remains unknown how the nsp3/4 pores assemble, if and how scission of DMVs from the ER occurs, and what fusogen accomplishes the complete separation of the two bilayers that delineate the DMV.

## Acknowledgments

We thank HMS Research Computing for resources supporting ColabFold, Talley Lambert (Nikon Imaging Center at HMS) for help with structural illumination microscopy, Maria Ericsson (Electron Microscopy Facility at HMS) for help with transmission EM, Lin Shao (Yale University) for CUDA-accelerated 3D-SIM reconstruction code, the Taplin Mass Spectrometry Facility at HMS for help with mass spectrometry, and the Center for Microscopy-Microanalysis and Information Processing at National University of Science and Technology Politehnica Bucharest for stimulated emission-depletion microscopy.

## Author contributions

A.C. and A.S. generated reagents and performed experiments. A.M.L. and A.C. performed ColabFold computing and A.M.L. performed modeling. S.G.S. and G.A.S. designed STED experiments and M.A. acquired and processed STED data. R.T.K. performed anti-viral assays with nucleoside analogs. K.H. participated in establishing hCoV-229E infection of host cells. T.d.A.M. participated in establishing the cell line expressing synthetic DMVs. T.J.M., P.A.S. and A.S. managed the team. A.C., A.S. and T.J.M. wrote the manuscript, with input from other authors.

## Data and material availability

All data are available in the main text or the supplementary materials

## Supplementary Materials

Materials and Methods Supplementary Text

Figs. S1 to S10 Tables S1

## Materials and methods

### DNA constructs

Plasmids carrying SARS-CoV2 and hCov-229E genes were obtained from the community depository of viral plasmids created by the Silver lab (Department of Systems Biology, Harvard Medical School), and were used as PCR templates for assembling various expression constructs. For bacterial expression, constructs were generated in the vectors pMal-c2 (NEB), pGEX-2tk (GE Healthcare) or pET30-HT7, a version of pET30 (EMD Millipore) for making N-terminal fusions to HaloTag7 (HT7, Promega). Fusions of the following proteins were expressed in bacteria: hCoV-229E N full-length, hCoV-229E nsp3 N-terminal fragment (nsp3-N, amino acids 300-703), hCoV-229E nsp4 C-terminal domain (nsp4-CTD, amino acids 370-480), hCoV-229E nsp7 full-length, hCoV-229E nsp8 full-length, hCoV-229E-nsp9 full-length, and an N-terminal fragment of human protein kinase R (PKR-N) comprising the two dsRNA-binding domains (amino acids 1-181). For constitutive expression in mammalian cells, the following constructs were subcloned into the pHAGE lentiviral vector [27]: SARS-CoV2 nsp4 tagged with 5 copies of the myc epitope at the N-terminus and mCherry at the C-terminus (myc-nsp4-mCherry), SARS-CoV2 nsp4 tagged with mCherry at the C-terminus (nsp4-mCherry), and nuclear-localized mCherry tagged with a HPC epitope at the N-terminus (NLS-HPC-mCherry). For tetracycline (Tet)-inducible expression in mammalian cells, a construct encoding N-terminally truncated SARS-CoV2 nsp3 (nsp3-DN, amino acids 1224-1945, starting with the b-coronavirus-specific bSM domain), tagged with eGFP at the N-terminus and HPC epitope at the C-terminus, was assembled in the lentiviral vector pLIX402 [28].

### Cell culture

Human embryonic kidney (HEK293) and HeLa cells were purchased from ATCC. Cells were grown in DMEM (Sigma) supplemented with 10% fetal bovine serum (Hyclone), 100IU/mL penicillin and 100µg/mL streptomycin (ThermoFisher), at 37°C and in an atmosphere of 5% CO_2_, in a humidified incubator. HEK293 cells were transiently transfected using Mirus TransIT-293 reagent (Mirus), following the protocol from the manufacturer.

### Stable cell lines

Constructs were built in third-generation lentiviral vectors and were used to produce lentiviruses by transient transfection of HEK293 cells. Stable cell lines were then generated by lentiviral transduction for 48hrs in the presence of polybrene (1µg/mL, Sigma), followed by selection with antibiotic (blasticidin, puromycin, or hygromycin), as described previously [29]. Expression of the construct of interest was confirmed by immunofluorescence and/or Western blotting. Single clones were isolated and characterized for the stable HEK293 cells expressing Tet-inducible eGFP-nsp3-DN-HPC together with constitutively-expressed nsp4-mCherry, used for assembling synthetic DMVs.

### Infection of cells with hCoV-229E

Media containing hCoV-229E was mixed with EMEM (ATCC) supplemented with 2% fetal bovine serum (Hyclone). The mix was then added to HeLa cells in a humidified incubator at 35°C and 5% CO2. The cells were fixed and analyzed by immunofluorescence at different times post-infection, to detect viral protein expression.

### Antibodies

Antibodies against hCoV-229E nucleocapsid (N) and non-structural proteins nsp3, nsp4, nsp7, nsp8, and nsp9 were raised in rabbits (Cocalico Biologicals) and were affinity-purified against antigen covalently immobilized on beads (Affi-Gel 10 or Affi-Gel 15, BioRad). Rabbit polyclonal antibodies against mCherry [27] and the mouse monoclonal antibody against human protein C (HPC) [29] were described before. Affinity-purified antibodies were fluorescently labeled by reaction with N-hydroxysuccinimidyl esters of various fluorophores (AlexaFluor488, AlexaFluor568, Cy-5), according to the manufacturers’ instructions. Antibodies were separated from unincorporated fluorophores on desalting columns (NAP-5 or PD-10, GE Healthcare).

The following antibodies were obtained commercially: mouse monoclonal anti-myc 9E10 (Sigma), mouse monoclonal anti-dsRNA J2 (Jena Bioscience), total rabbit IgG (Jackson Immunoresearch), mouse monoclonal anti-α-tubulin DM1α (Santa Cruz Biotechnology), CF488-conjugated goat anti-rabbit secondary (Biotium), CF568-conjugated goat anti-rabbit secondary (Biotium), AlexaFluor594-conjugated goat anti-rabbit secondary (ThermoFisher), AlexaFluor647-conjugated goat anti-mouse secondary (ThermoFisher), Abberior STAR RED 647-conjugated goat anti-rabbit secondary (Abberior).

### Reagents

The following reagents were purchased: Hoechst 33342 (Sigma), ProLong Diamond mountant (ThermoFisher), methanol-free formaldehyde (ThermoFisher), Tosylactivated Dynabeads (ThermoFisher), Protein-A agarose (Sigma), Streptavidin agarose (ThermoFisher), digitonin (Calbiochem), 5’-ethynyl-uridine (Cayman Chemical), actinomycin D (Cayman Chemical), biotin-azide (MedChem Express), N6-[2-(4-aminophenyl)ethyl]-adenosine (MedChem Express), N6-(4-hydroxybenzyl)-adenosine (MedChem Express), N6-benzyl-adenosine (MedChem Express), N6-methyl-adenosine (MedChem Express), N6-dimethyl-adenosine (MedChem Express), N6-propargyl-adenosine (Jena Bioscience), CF647-conjugated streptavidin (Biotium), aminoallyl-UTP-PEG_5_-AlexaFluor647 (Jena Bioscience), AlexaFluor488-NHS ester (ThermoFisher), AlexaFluor568-NHS ester (ThermoFisher), Cy5-NHS ester (MedChem Express), HaloTag succinimidyl ester (O2) ligand (Promega), HaloTag-Oregon Green ligand (Promega), HaloTag-TMR ligand (Promega), HaloTag-biotin ligand (Promega), horseradish peroxidase (HRP, ThermoFisher).

### Protein expression and purification from bacteria

Fusion proteins were expressed in BL21(DE3)pLysS E.coli (EMD Millipore). The proteins were tagged at the N-terminus with his_6_-HT7 (pET30-HT7 vector), maltose-binding protein (MBP) (pMal-c2 vector) or glutathione-S-transferase (GST) (pGEX-2tk vector). Bacterial pellets were lysed in lysis buffer (50mM Tris pH 7.5, 150mM NaCl, 1% Triton X-100, 1mM PMSF). In the case of his_6_-HT7 fusions, the lysis buffer was supplemented with 25mM imidazole pH 7.5.

Lysates were clarified by centrifugation and the recombinant protein was purified according to the manufacturers’ instructions, using Ni-NTA agarose (Qiagen), amylose resin (NEB), or glutathione agarose (GE Healthcare). Eluted protein was dialyzed against HEPES-buffered saline (HBS, 10mM Na-HEPES pH 7.5, 150mM NaCl) overnight at 4°C, concentrated using Amicon centrifugal filters (Sigma), frozen in liquid nitrogen and stored at −80°C. The probe for dsRNA detection consisted of his_6_HT7-tagged hPKR-N. The following proteins were used for rabbit immunizations: MBP-229E-N, MBP-229E-nsp3-N, GST-229E-nsp4-CTD, his_6_HT7-229E-nsp7, MBP-229E-nsp8, and his_6_HT7-229E-nsp9. The following proteins were used to generate affinity matrices for antibody purification from immune sera: GST-229E-N, GST-229E-nsp3-N, GST-229E-nsp4-CTD, his_6_HT7-229E-nsp7, his_6_HT7-229E-nsp8, and MBP-229E-nsp9.

### Double-stranded RNA (dsRNA) probes

Viral dsRNA was detected using purified recombinant his_6_Halo-hPKR-N labeled with a fluorophore, biotin, or HRP. Fluorescently-and biotin-labeled his_6_Halo-hPKR-N were generated by incubating the recombinant protein with HaloTag-fluorophore or HaloTag-biotin ligands, according to the manufacturer’s instructions. Excess HaloTag ligand was then removed by purification on desalting NAP-5 or PD-10 columns (GE Healthcare). To generate the HRP-his_6_Halo-hPKR-N conjugate, purified HRP was first modified with HaloTag succinimidyl ester (O2) ligand and was then reacted with purified his_6_Halo-hPKR-N; we estimated that a 1:1 molar ratio between the proteins resulted in >90% modification of his_6_Halo-hPKR-N. An immunofluorescence protocol (see below) was used to stain fixed virus-infected cells with various dsRNA probes.

### Immunofluorescence

HEK293 cells expressing genes of interest or HeLa cells infected with hCoV-229E were fixed with 2% formaldehyde in phosphate-buffered saline (PBS), for 10min at room temperature. After washing with PBS, the cells were permeabilized with 40µM digitonin for 3-5min, or with 0.2% Triton X-100 for 10min. Cells were counterstained with Hoechst 33342 (1µg/mL) and blocked with blocking solution (3% BSA in PBS). For indirect immunofluorescence, the cells were then incubated with unlabeled primary antibodies diluted in blocking solution, overnight at 4°C. After washing with PBS, the cells were incubated with labeled secondary antibodies in blocking buffer, for 1h at room temperature. The cells were washed in PBS, were mounted in Prolong Diamond mounting media, and were imaged by fluorescence microscopy. For direct immunofluorescence, cells were stained with labeled primary antibodies diluted in blocking buffer, overnight at 4°C. For detection of dsRNA, virus-infected cells were incubated with fluorescently-conjugated his_6_Halo-hPKR-N diluted in blocking buffer, for 15min at room temperature.

### Detection of viral RNA synthesis in cells by click chemistry

To label viral RNA with 5’-ethynyl uridine (EU), hCoV-229E-infected HeLa cells were pre-incubated with 10µM actinomycin D for 1hr (to suppress host cell transcription). The cells were then incubated with 1mM EU in the continued presence of actinomycin D, for 45min, followed by fixation with 2% formaldehyde in PBS for 15min at room temperature. The cells were permeabilized with Tris-buffered saline (TBS) with either 40µM digitonin or with 0.2% Triton X-100 (TBST), washed with TBS, and then stained with 10µM biotin-azide in click buffer (100mM Tris pH 8.5, 1mM CuSO_4_, 100mM ascorbic acid) for 30min. The cells were then washed with TBS and stained with fluorescent streptavidin conjugates diluted in blocking buffer, followed by fluorescence microscopy. For testing differential digestion of viral RNA by nucleases, EU-labeled cells fixed and permeabilized as above were incubated in 1×CutSmart buffer (NEB) with 4ng/µL RNase A or 5U/µL benzonase, for 1h at 37°C. Cells were then washed thoroughly with TBS and incorporated EU was detected by click reaction as above.

### Immunoprecipitation

To immunopurify DMVs, virus-infected HeLa cells were resuspended in hypotonic lysis buffer (10mM Na-HEPES pH 7.6, 10mM Na acetate, 1.5mM MgCl_2_) and were lysed on ice using a Dounce homogenizer (30 strokes). The lysate was centrifuged for 10min at 650g and 4°C for 10min, the nuclear pellet was resuspended in HBS, and was incubated with anti-nsp3 antibodies immobilized on magnetic beads, for 2hrs at 4°C. The beads were washed 3 times with HBS, and precipitated proteins or RNAs were analyzed by immunoblotting or qRT-PCR, respectively. To immunoprecipitate viral proteins, virus-infected HeLa cells or nuclear pellets thereof (see above) were lysed in HBS with 1% Triton X-100. The lysate was clarified by centrifugation for 20 min at 14,000g and 4°C, and was incubated with anti-nsp3, anti-nsp8, or total rabbit IgG (negative control) antibodies immobilized on magnetic beads, for 2hrs at 4°C. The beads were washed 3 times with HBS with 1% TX-100, and co-purified viral proteins and RNAs were analyzed as above for immunopurified DMVs.

### Protein electrophoresis and immunoblotting

Protein samples were mixed with SDS sample buffer supplemented with 50mM DTT, boiled and then subjected to separation by SDS-PAGE (4-20% Criterion TGX Precast Midi Gels, BioRad). Separated proteins were visualized by staining with Coomassie Blue R250 (G-Biosciences).

Alternatively, the separated proteins were transferred to polyvinylidene difluoride (PVDF) membranes (BioRad), which were rinsed with TBST, blocked with 5% non-fat milk in TBST, and incubated with primary antibodies in TBST with milk, overnight at 4°C. The membrane was then washed with TBST and incubated with HRP-conjugated secondary antibodies for 1h at room temperature. The membrane was washed again, incubated for 1min with chemiluminescent substrate (Revvity), and then imaged (Amersham ImageQuant 800, Cytiva).

### Isolation of synthetic DMVs by sucrose gradient centrifugation

Sucrose gradients (20%-60%) were prepared in centrifugation buffer (10mM Na-HEPES pH 7.6, 75mM NaCl), in 14×95mm polypropylene tube (Beckman), using Gradient Master (BIOCOMP). HEK293 cells expressing synthetic DMVs were harvested and resuspended in ice-cold buffer containing 10mM HEPES pH 7.6, 1.5mM MgCl_2_, 10mM Na acetate. The cells were lysed in a Dounce homogenizer on ice, with 15 strokes. Lysates were centrifuged for 15min at 500g and 4°C. The supernatant was collected, adjusted to 75mM NaCl and 5% sucrose, and was layered on top of the sucrose gradient. The sucrose gradients were centrifuged for 4h at 36000rpm and 4°C, in a SW-40 rotor (Beckman). Gradient fractions were collected using a gradient fractionation system (BRANDEL) and were analyzed by SDS-PAGE or immunoblotting. Fractions enriched in eGFP-nsp3-DN-HPC and nsp4-mCherry were pooled and subjected to immunoprecipitation with anti-HPC antibody covalently immobilized on magnetic beads (Tosylactivated Dynabeads, ThermoFisher). The beads were washed 3 times with IP buffer and fixed by a fixation buffer (described in EM method section) for electron microscopy analysis.

### Proximity labeling assay and denaturing co-IP

HeLa cells infected with hCoV-229E were fixed with 1% formaldehyde in PBS, for 10min at room temperature. Cells were washed in PBS, permeabilized, and subjected to immunostaining with primary antibodies against nsp4, nsp7 or nsp8, and then with HRP-conjugated goat anti-rabbit secondary antibody. After washing with PBS, cells were incubated with 5µM biotin-tyramide (MiliporeSigma) in HRP staining buffer (100mM Tris pH 7.5, 150mM NaCl, 0.1% Tween-20, 0.003% H_2_O_2_), for 15min at room temperature. Cells were washed with TBS, collected and incubated for 1h at 65°C in 50mM Tris pH 7.5, 150mM NaCl, 2.5% SDS. The samples were centrifuged for 15min at 14000g, the supernatant was diluted with TBS to 0.5% SDS, and was incubated with streptavidin-agarose for 2hrs at room temperature. The beads were washed with TBS+0.5% SDS, and the bound protein was eluted with 62.5mM Tris pH 6.8, 2%SDS, 10% glycerol, 1mg/mL bromophenol blue at 65°C for 15min and was analyzed by SDS-PAGE and immunoblotting.

### Run-off viral RNA synthesis

Virus-infected HeLa cells were rinsed in ice-cold PBS, incubated with 40µM digitonin for 3-5min on ice, and then washed thoroughly with run-off buffer [10mM K-HEPES pH 7.6, 50mM KCl, 2mM MgCl_2_, 200mM sucrose, protease inhibitor cocktail (MiliporeSigma)]. The cells were incubated afterwards with run-off buffer supplemented with NTPs (250µM each) and aminoallyl-UTP-PEG_5_-AlexaFluor647 (50µM), for 1h at 35°C. Following fixation, the cells were washed and imaged by fluorescence microscopy.

### Viral RNA synthesis on beads

Nuclear pellets from hypotonically lysed hCoV-229E-infected HeLa cells were solubilized in TBST and were incubated with anti-nsp3, anti-nsp8 or rabbit total IgG (negative control) antibodies immobilized on magnetic beads. After washing, the beads were incubated with PBS supplemented with 0.02% TX-100, 2mM MgCl_2_, 250µM of each NTP, and 100µM aminoallyl-UTP-PEG_5_-AlexaFluor647, for 1h at 35°C. The beads were washed and imaged by fluorescence microscopy.

### Quantitative RT-PCR

Total RNA co-immunoprecipitated with viral proteins was isolated using Trizol (ThermoFisher), according to the manufacturer’s protocol. Purified RNA was reverse transcribed with Luna Superscript Mix (NEB). RNA transcripts of viral and host genes were measured by quantitative PCR using primers listed in Table 2 and Power SYBR Green kit (ThermoFisher), on a QuantStudio7 Pro Real-time PCR System (ThermoFisher).

### Mass spectrometric analysis

Co-immunoprecipitated proteins were eluted from beads with SDS sample buffer (62.5mM Tris pH 6.8, 2%SDS, 10% glycerol, 1mg/mL bromophenol blue), for 15min at room temperature. The eluate was supplemented with DTT to 50mM and was incubated for 10min at 60°C. The samples were loaded onto a 4-20% Criterion TGX Precast Midi Gel (BioRad) and were separated until the dye front migrated ∼1.5cm. The gel was fixed in 50% ethanol and 10% acetic acid for 30min and was stained with Coomassie Blue R250 (G-Biosciences). The portions of the gel containing the samples were excised and the gel pieces were subjected to in-gel trypsin digestion [30].

Tryptic peptides extracted from the gel were then dried and stored at 4°C until mass spectrometric analysis. The peptide samples were separated on a nano-scale reverse-phase HPLC capillary column (100 µm inner diameter and 30 cm length) packed with 2.6 µm diameter spherical C18 silica beads, using a linear gradient of acetonitrile [31]. Eluting peptides were subjected to electrospray ionization and then introduced into an Orbitrap Exploris480 mass spectrometer (ThermoFisher). Peptides were detected, isolated, and fragmented, and the resulting fragmentation pattern was used to determine peptide sequences using Sequest software (ThermoFisher) [32].

### Microscopic imaging

Cells were grown on glass coverslips (#1.5, Electron Microscopy Sciences), were processed for immunofluorescence, and were imaged by spinning-disc confocal microscopy, 3D-structural illumination microscopy (3D-SIM), or stimulated emission depletion (STED) microscopy.

Spinning-disk confocal imaging was performed on a Nikon Ti-E inverted microscope equipped with a Yokagawa CSU-X1 spinning disk unit and ORCA-Fusion BT sCOMS camera, and controlled with the Nikon NIS-Elements software. Images were acquired using either a Nikon PlanFluor 40× 1.30NA oil objective or a Nikon PlanApo 100× 1.45NA oil objective. For deconvolution of confocal image stacks acquired with the 100× objective, a theoretical point spread function was generated with the PSF Generator plugin using the Born & Wolf 3D Optical Model, with a voxel depth of 0.3µm and 1.45NA. Afterwards, the deconvolution was done with the DeconvolutionLab2 plugin utilizing a Richardson-Lucy algorithm with 10 iterations.

3D-SIM imaging was performed on an OMX V4 Blaze microscope (GE Healthcare) equipped with three PCO.edge sCMOS cameras, and 488 nm and 568 nm laser lines. Images were acquired with a 0.125 mm step size without binning, using a 60x 1.42NA PlanApochromat objective (Olympus). For each z-section, 15 raw images (three rotations with five phases each) were acquired. Spherical aberration was minimized using immersion oil matching [33]. Super-resolution images were computationally reconstructed from the raw data sets with a channel-specific, measured optical transfer function and a Wiener filter constant of 0.001, using CUDA-accelerated 3D-SIM reconstruction code based on Gustafsson et al. (2008) [34]. TetraSpeck beads (Thermo Fisher) or a nano-grid control slide (GE Healthcare) were used to measure axial and lateral chromatic misregistration, and experimental data sets were registered using the imwarp function in MATLAB (MathWorks).

Stimulated Emission Depletion (STED) imaging was performed on an Abberior Instruments Expert Line system working in a time-gated configuration, implemented on an inverted Olympus IX83 microscope. Images were acquired with an Olympus UPlanSApo 100× 1.4NA objective using 1.518 RI immersion oil. AlexaFluor594 and Abberior STAR RED signals were acquired by using 561 nm and 640 nm laser lines for excitation, respectively. For both channels, we used a 775 nm STED beam for depletion. For the AlexaFluor594 channel, fluorescence was detected in the range of 595-665 nm, and for the Aberrior STAR RED channel in the 650-750nm range. For a region of interest, Z-stacks were acquired for a range of a few microns (depending on the size of the imaged structures), using a z-step of 100nm. For deconvolution purposes, xy scanning was oversampled by using a 10 nm pixel size. STED images were deconvolved with Huygens Professional software (SVI, The Netherlands), using the express wizard operating in standard profile. For deconvolution parameters, a STED saturation factor of 30 and a STED immunity fraction of 10% were used.

### Image analysis

For quantifying the number and intensity of nsp and dsRNA puncta, images were analyzed using the “analyze particle” function in ImageJ. All images acquired for an experiment were subjected to the same thresholding. For profiling the signal intensity distribution of puncta in an image, mean intensities were assigned to 300 bins in all experimental groups. Fluorescent intensity line scans were generated using the “plot profile” function, and line scans of different channels were merged into one graph. Colocalization analysis was performed using the coloc-2 plugin, and a Pearson coefficient was used assess the degree of colocalization. Three non-overlapping fields of view were analyzed for each experimental condition, and each condition was repeated 3 times, on different days.

### Electron microscopy

For visualizing DMVs in cells, HEK293 cells expressing synthetic DMVs or HeLa cells infected with hCoV-229E were grown as monolayers on Aclar coverslips (Electron Microscopy Science). The cells were fixed with 1.25% formaldehyde, 2.5 % glutaraldehyde and 0.03% picric acid in 0.1M pH 7.4 sodium cacodylate buffer, for 2hrs at room temperature. The coverslips were then washed in cacodylate buffer and were post-fixed in 1% osmium tetroxide/1.5% potassium ferrocyanide for 1 hour, followed by washes with water and 50mM maleate buffer pH 5.15 (MB). After incubation with 1% uranyl acetate in MB for 1hr, the cells were washed with MB and water, followed by stepwise dehydration to 100% ethanol. After carefully removing the ethanol, the cells were layered with a drop of TAAB Epon (TAAB Laboratories Equipment Ltd) and covered with another sheet of Aclar, followed by polymerization at 60°C for 48 hrs. The top Aclar was then peeled off, and a small area (∼1mm) of the embedded cells was cut out with a razor blade and remounted on an Epon block. Ultrathin sections (80nm) were cut on a Reichert Ultracut-S microtome, picked up on copper grids stained with lead citrate, and imaged on a Tecnai G2 Spirit BioTWIN transmission electron microscope equipped with an AMT NanoSprint43-MKII (43 megapixel) CCD camera.

### Data quantification and statistical analysis

Statistical information for each experiment is provided in the figure legends. For all image analysis, at least three non-overlapping fields of view were acquired per condition, and each experiment was repeated at least three times, on different days. All biochemistry experiments (including synthetic DMV isolation and detection, immunoprecipitation of viral proteins, dsRNA affinity isolation, proximity labeling, quantitative PCR, *in-vitro* viral RNA synthesis) were repeated independently at least three times. Mass spectrometric and electron microscopy experiments were performed at least twice, on different days.

---

**Fig. S1.**
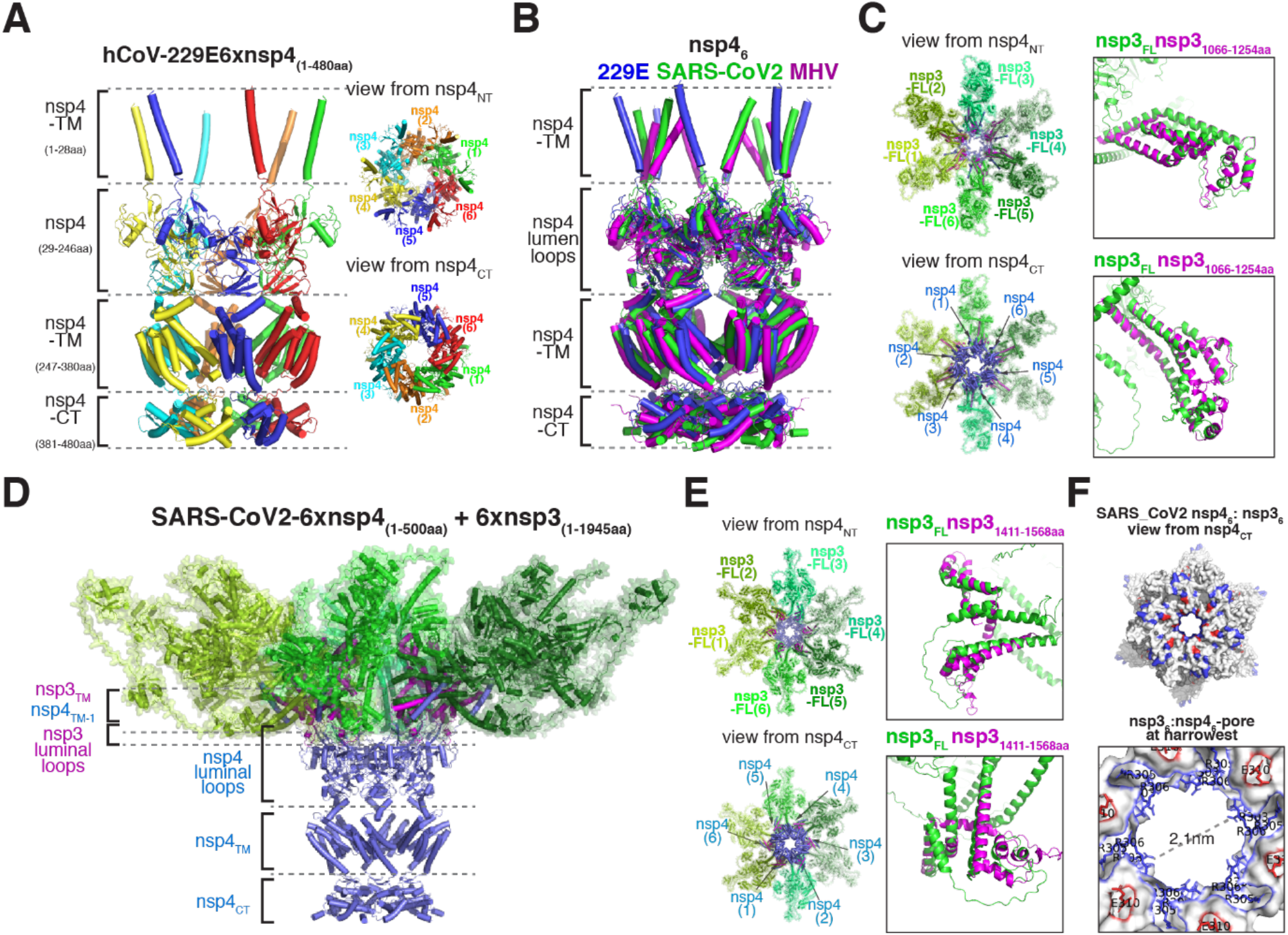
Predicted nsp3-nsp4 and replicase subunits-nsp4 complexes. **(A)** Predicted structure of a six-fold symmetric complex of nsp4 from hCoV-229E, seen from the side (left) and from the top and bottom (right). TM = transmembrane domains; NT = N-terminus; CT = C-terminus. **(B)** As in (A), but including hCoV-229E (blue), SARS-CoV2 (green), and MHV (purple). **(C)** Predicted structure of a six-fold symmetric complex between hCoV-229E nsp3 (residues 1066-1254) and full-length nsp4. Left: top and bottom views of ribbon diagrams; right: close-up view of the alignment between hCoV-229E full length nsp3 (green) and nsp3 (1066-1254aa) (purple). **(D)** Predicted structure of a six-fold symmetric complex between SARS-CoV2-nsp3 (residues 1411-1568aa) and full-length nsp4, including the predicted structure of full-length SARS-CoV2-nsp3, seen from the side. **(E)** As in (D), but showing top and bottom views of the complex, respectively. Left: ribbon diagram; right: close-up view of the alignment between SARS-CoV2 full length nsp3 (green) and nsp3 (1411-1568aa) (purple). **(F)** As in (D), Top: molecular surface showing positive (blue) and negative (red) residues viewing from nsp4_CT_; bottom: predicted narrowest portion of the pore formed by SARS-CoV2 nsp4 in complex with nsp3. The diameter of the constriction and the conserved residues delineating it are indicated.

**Fig. S2.**
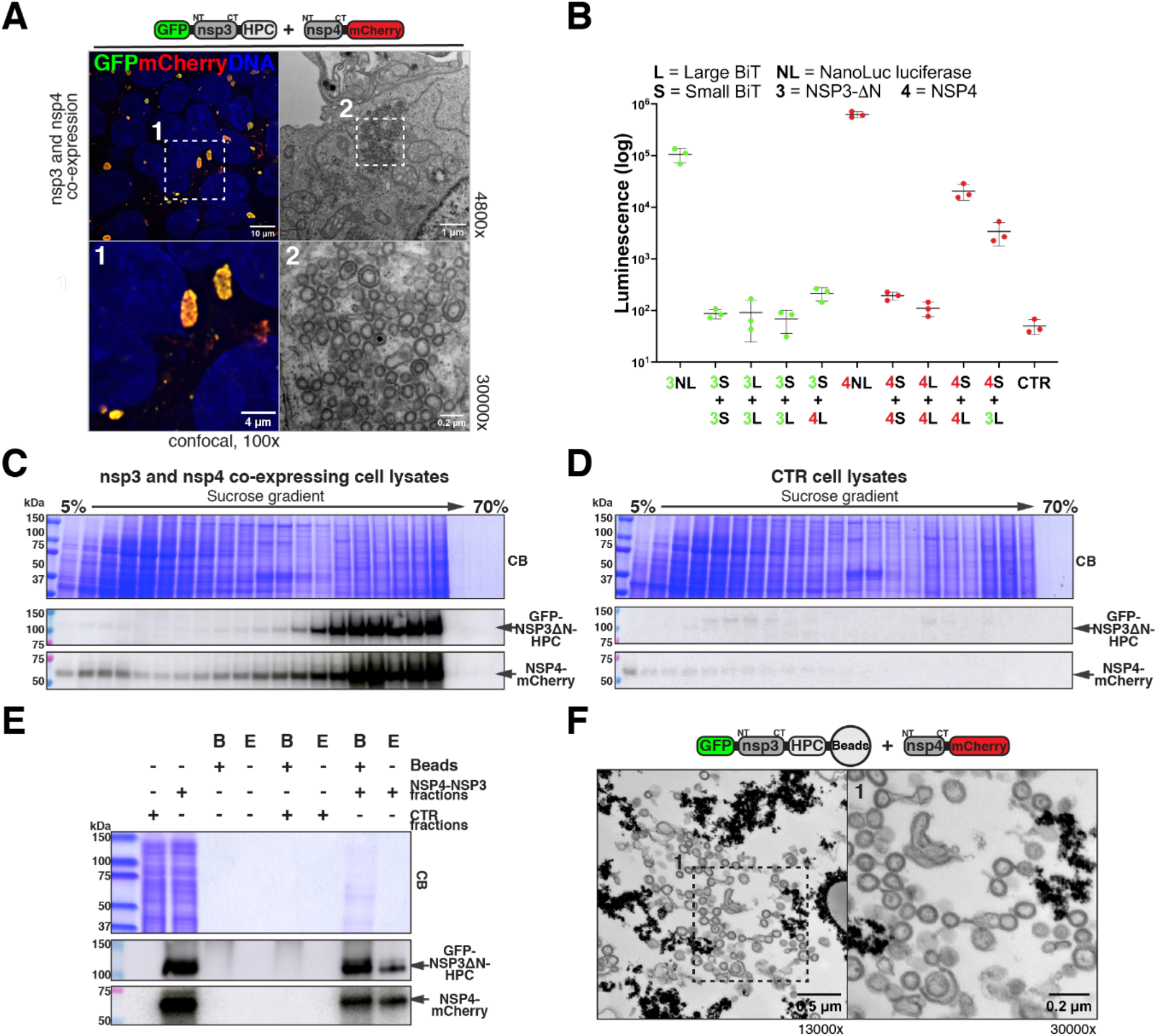
DMV reconstitution by nsp3-nsp4 co-expression and DMV affinity isolation. **(A)** Fixed HEK293 cells expressing eGFP-nsp3ΔN-HPC and nsp4-mCherry were imaged by confocal microscopy (100×). Nsp3 and nsp4 colocalize. **(B)** Cells expressing nsp3-NanoLuc (NL), nsp4-NanoLuc (NL), or co-expressing a combination (as indicated in the graph) from the following constructs: nsp3ΔN-LargeBit (3L), nsp3ΔN-SmallBit(3S), nsp4-LargeBit(4L), nsp4-SmallBit(4S), were lysed and assayed for luciferase activity. Robust luciferase activity is observed when 3S and 4L, or 4S and 4L are co-expressed, indicating nsp3 interacts with nsp4 and nsp4 interacts with itself. **(C)** HEK293 cells co-expressing GFP-nsp3ΔN-HPC and nsp4-mCherry were subjected to hypotonic lysis (detergent-free), and lysates were separated by ultra-centrifugation on a sucrose gradient (5%-70%). Gradient fractions were analyzed by SDS-PAGE and immunoblotting with anti-HPC or anti-mCherry antibodies. Nsp3 and nsp4 are enriched in the same gradient fractions. **(D)** As in (C), but using control HEK293 cells. **(E)** As in (C), but nsp3-and nsp4-enriched gradient fractions were subjected to immunoprecipitation using anti-HPC antibodies. Precipitated material was eluted from beads with the HPC peptide. Material on beads (B) and in the eluate (E) was analyzed by SDS-PAGE and immunoblotting with anti-HPC or anti-mCherry antibodies. **(F)** As in (E), but the beads were imaged by TEM (magnification 13000×, left; 30000×, right). Synthetic DMVs are isolated on beads.

**Fig. S3.**
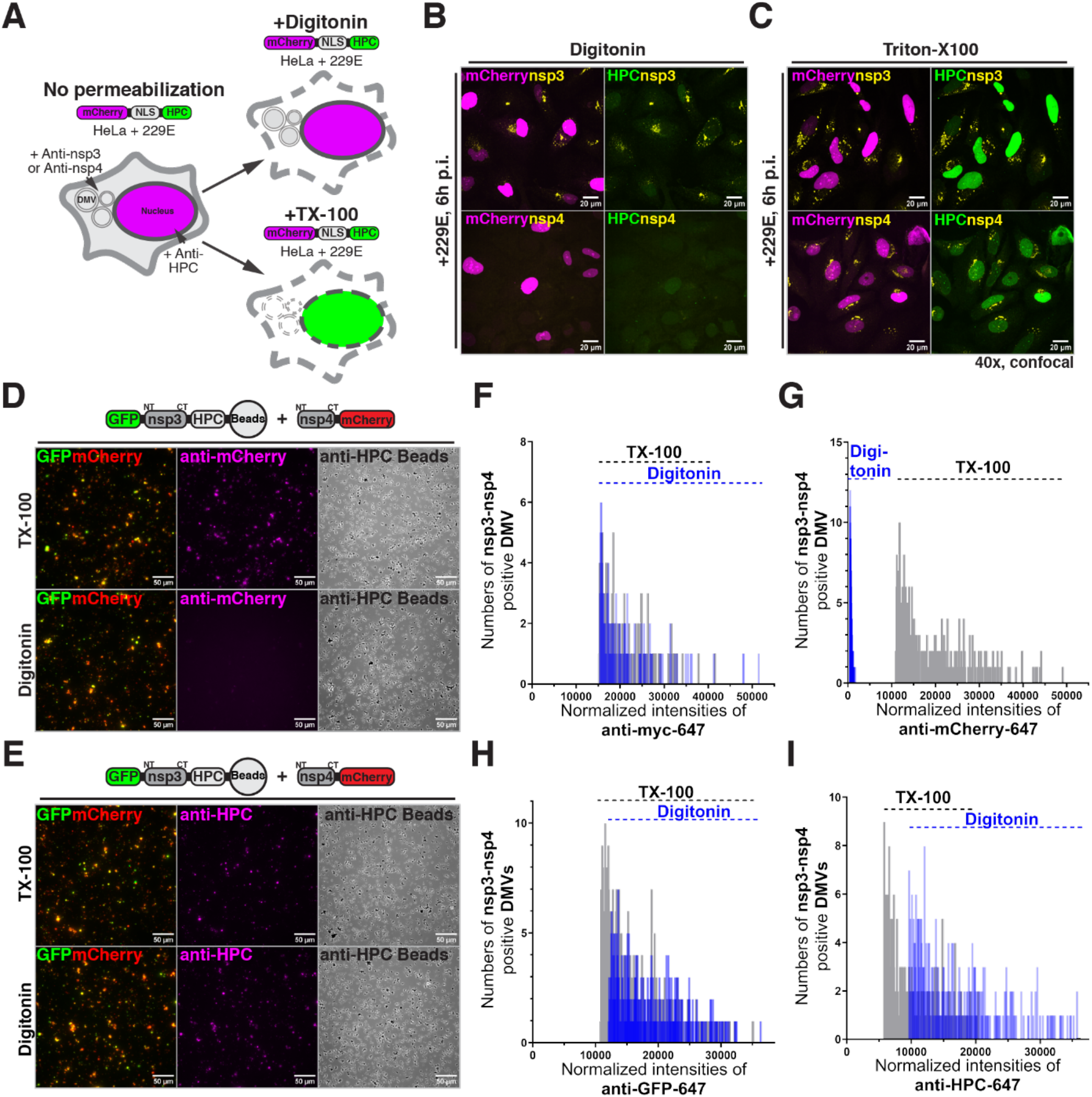
Differential accessibility of assays for nsp3 and nsp4. **(A)** Schematic of differential antibody accessibility assay. HeLa cells expressing HPC-tagged NLS-mCherry were infected with hCoV-229E. Cells were fixed, differentially permeabilized and stained with fluorescent anti-HPC antibodies. **(B)** As in (A), but cells permeabilized with digitonin and co-stained with anti-nsp3-NT and anti-HPC antibodies (top panel) or with anti-nsp4-CT and anti-HPC antibodies (bottom panel). Only the cytoplasm-facing nsp3-NT is detected by antibodies. **(C)** As in (B), but with TX-100 permeabilization. HPC-NLS-mCherry, nsp3 and nsp4 are all detected. **(D)** Synthetic DMV were isolated from HEK293 cells co-expressing eGFP-nsp3ΔN-HPC and nsp4-mCherry on anti-HPC beads (described in Fig. S2C-F). The beads were fixed, differentially permeabilized and stained with anti-mCherry antibodies. The mCherry tag is detected with TX-100 but not with digitonin. **(E)** As in (D), but with anti-HPC antibodies. The HPC tag nsp3 C-terminus is detected with both digitonin and TX-100. **(F-I)** Fixed HEK293 cells co-expressing eGFP-nsp3ΔN-HPC and myc-nsp4-mCherry were differentially permeabilized and stained with indicated antibodies. Measured DMV intensities were assigned into 300 bins and the number of DMVs in each bin were quantified. EGFP-, HPC-, or myc-positive DMVs have similar intensity profile between digitonin (blue) and TX-100 (grey) permeabilization, while mCherry-positive DMVs have much weaker intensity profile with digitonin.

**Fig. S4.**
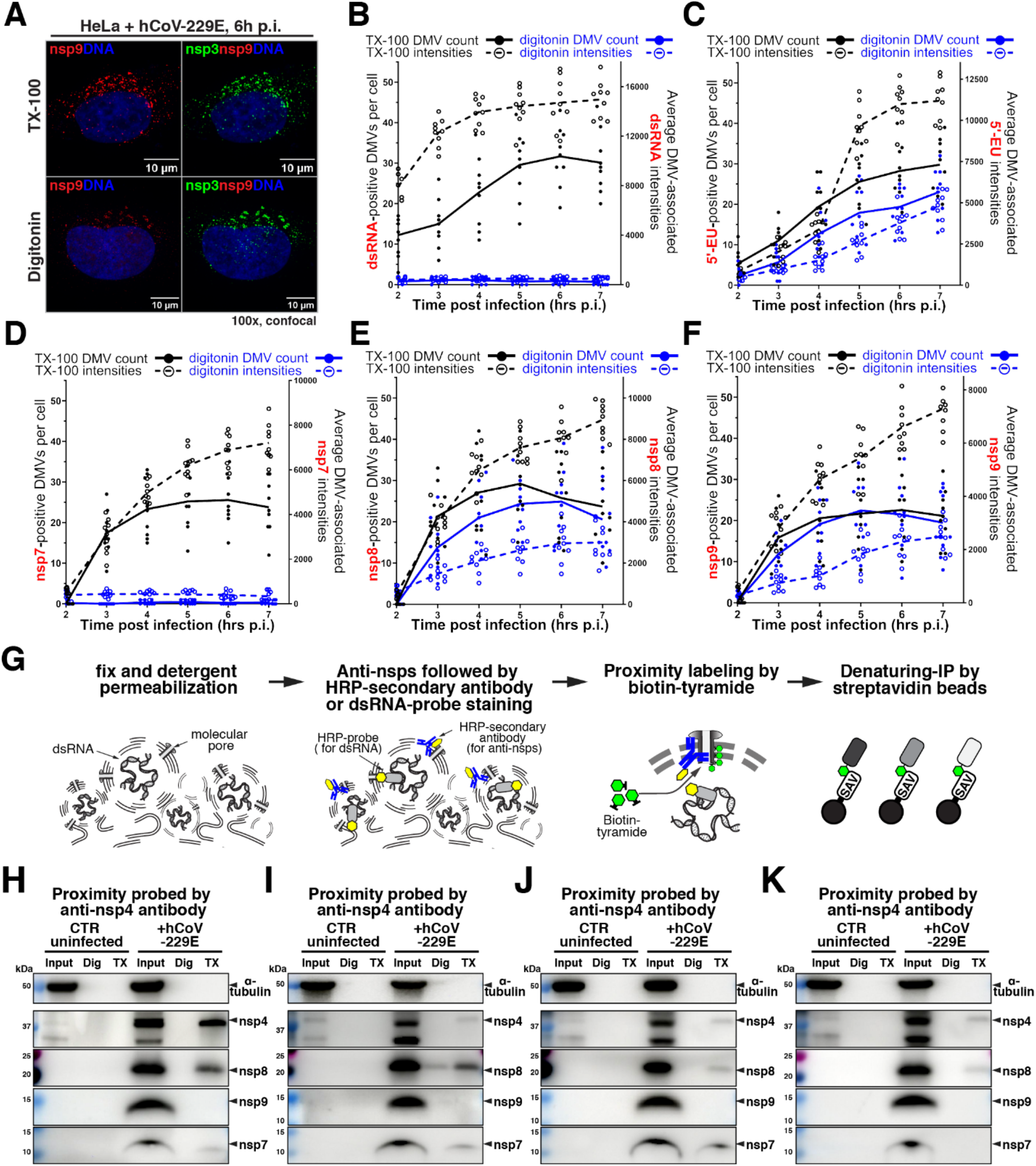
Replicase subunits and nsp4 localize inside DMVs at early infection stages. **(A)** hCoV-229E-infected HeLa cells were assayed for differential accessibility using a mix of fluorescently labeled anti-nsp3 and anti-nsp9 antibodies. Nsp9 is weakly accessible with digitonin permeabilization, in contrast to nsp3. **(B-F)** hCoV-229E-infected HeLa cells were assayed for differential accessibility using the dsRNA probe (B), or with click chemistry detection of EU incorporated at DMVs (C), or anti-nsp7 antibodies (D), or anti-nsp8 antibodies (E), or anti-nsp9 antibodies (F) at different post infection time points. Three non-overlapping fields of view were analyzed per condition and each condition was repeated 3 times. (*N*=3, n=9). **(G)** Schematic of proximity detection of viral nsps and dsRNA. HeLa cells are infected with hCoV-229E, lightly fixed and permeabilized, and stained with antibodies against nsp4, nsp7 or nsp8, or HRP-conjugated dsRNA probe. Primary antibodies against nsps are stained with HRP-conjugated secondary antibodies. The cells are then reacted with biotin-tyramide, and biotinylated viral nsps are affinity purified on streptavidin beads and analyzed. **(H-J)** As in (G), cells were stained with anti-nsp4 (H), or anti-nsp7 antibodies (I), or HRP-conjugated dsRNA probe (J) followed by proximity detection. Affinity purified biotinylated material was analyzed by SDS-PAGE and immunoblotting with antibodies against nsp4, nsp7 and nsp8.

**Fig. S5.**
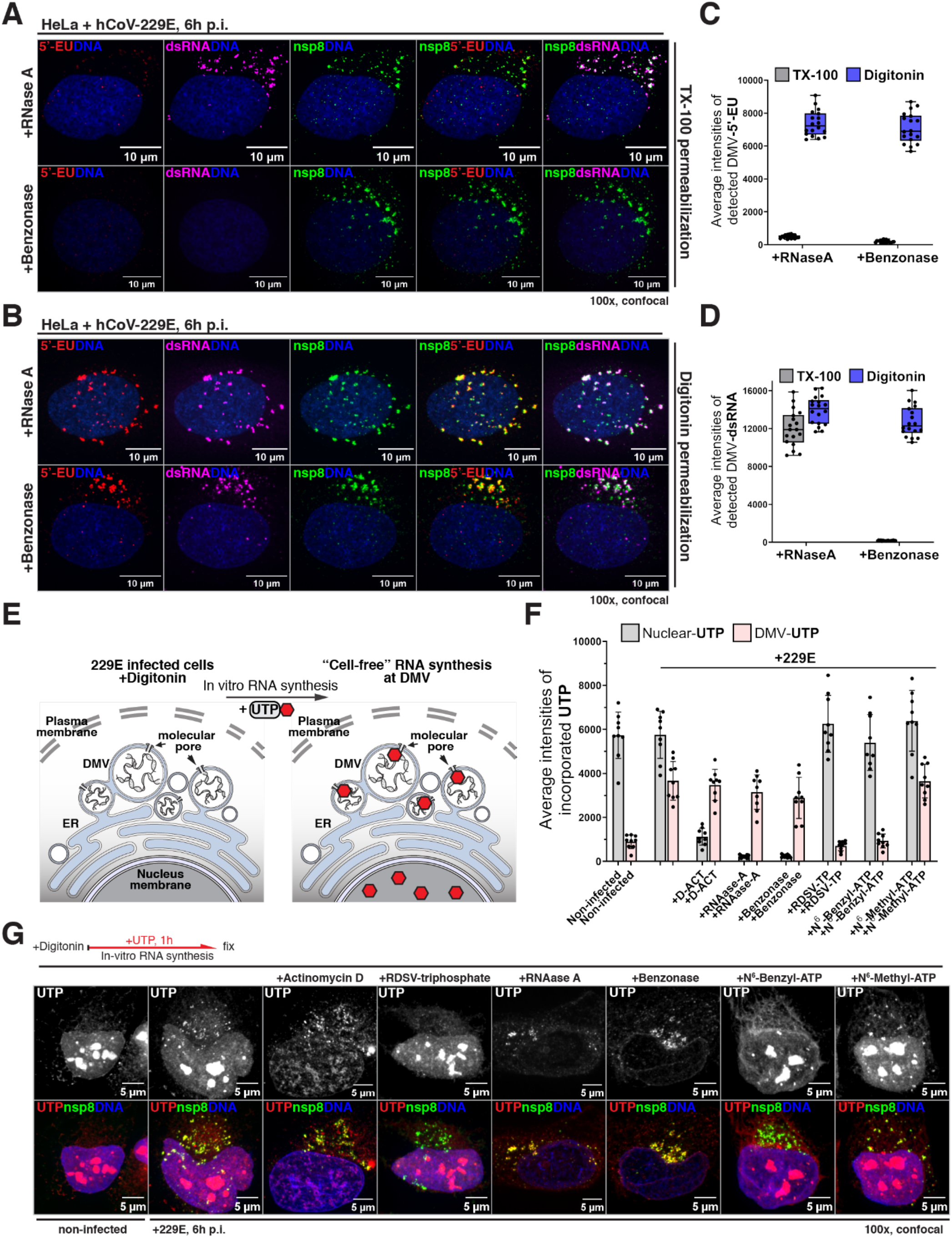
Nascent viral mRNA is synthesized within DMVs and then spreads to cytoplasm. **(A and B)** hCoV-229E-infected HeLa cells were labeled with EU, fixed and permeabilized by TX-100 (A) or digitonin (B). The cells were digested with enzymes and reacted by click reaction with biotin-azide, followed by staining with fluorescent streptavidin, dsRNA probe and anti-nsp8 antibodies. **(C and D)** Quantification of average EU or dsRNA intensities at DMV under different conditions in (A and B). EU-labeled RNA is digested by RNase A with TX-100, but resistant to RNase A with digitonin permeabilization. DsRNA is only digested by benzonase with TX-100 permeabilization. Thus, nascent viral mRNA spreads from inside DMVs to the cytoplasm, but dsRNA resides strictly inside DMVs. **(E)** Schematic of run-off RNA synthesis assays. hCoV-229E-infected HeLa cells were permeabilized with digitonin and incubated with a mix of NTPs and fluorescent UTP. Cells were then fixed and stained with anti-nsp8 antibodies. **(F and G)** As in (E), but RNA synthesis was performed in the presence of indicated compounds or enzymes. Fluorescent UTP intensity in DMVs and nucleus was measured, in three non-overlapping fields of view per condition; each condition was repeated 3 times. (*N*=3, n=9). UTP incorporation into DMVs is inhibited by co-incubation with remdesivir triphosphate, and is resistant to RNase digestion.

**Fig. S6.**
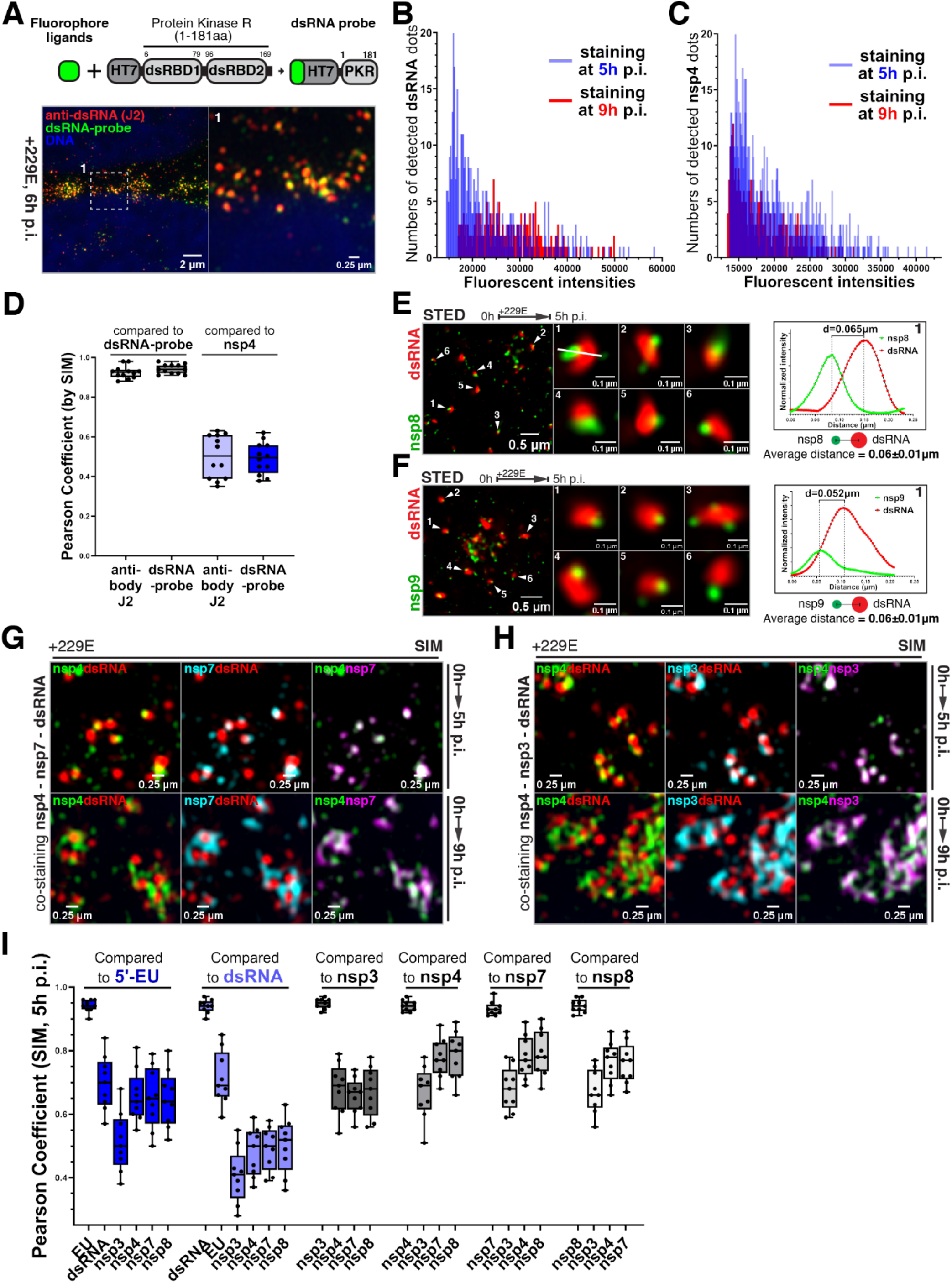
High resolution imaging of dsRNA, nsp8 and nsp9. **(A)** Top: schematic of the dsRNA-specific probe. Bottom: hCoV-229E-infected HeLa cells were fixed and co-stained with the dsRNA probe and anti-dsRNA antibodies (J2). **(B and C)** As in (A), but cells co-stained with dsRNA-probe and anti-nsp4 antibodies at different post infection time points. Average intensities of dsRNA and nsp4 signals at DMVs were assigned to 300 bins, and the number of DMVs in each bin were quantified. The dsRNA intensity profile (but not nsp4) shifts from a peak at 5 hours to an even distribution at 9 hours. **(D)** As in (A-C), showing Pearson coefficients for each indicated co-detected pair. Four non-overlapping fields of view were analyzed per condition and each condition was repeated 3 times. (*N*=3, n=12). **(E, F)** As in (A), but cells co-stained with anti-nsp8 (E) or anti-nsp9 (F) antibodies and dsRNA probe, then imaged by stimulated emission depletion microscopy (STED). Middle panels show zoomed-in views of the DMVs indicated in the image on the left. The white line in middle panel 1 was used for linescan analysis (right panel), to measure distance between nsp8 and dsRNA signals. **(G and H)** As in (B and C), but co-stained with anti-nsp4, anti-nsp7, and dsRNA probe (G); or anti-nsp4, anti-nsp3 and dsRNA probe (H). **(I)** hCoV-229E-infected HeLa cells were fixed, permeabilized and co-stained with all possible double combinations of nsp3, nsp4, nsp7, nsp8, dsRNA and nascent RNA (EU). The cells were imaged by SIM and Pearson coefficients were quantified for all pairs. Three non-overlapping fields were analyzed for each co-detected pair, and each staining condition was repeated 3 times (*N*=3, n=9). All nsps except nsp3 strongly co-localize with EU-labeled RNA, while showing less co-localization with dsRNA. Nsp3 is an outlier, localizing away from all the other markers above.

**Fig. S7.**
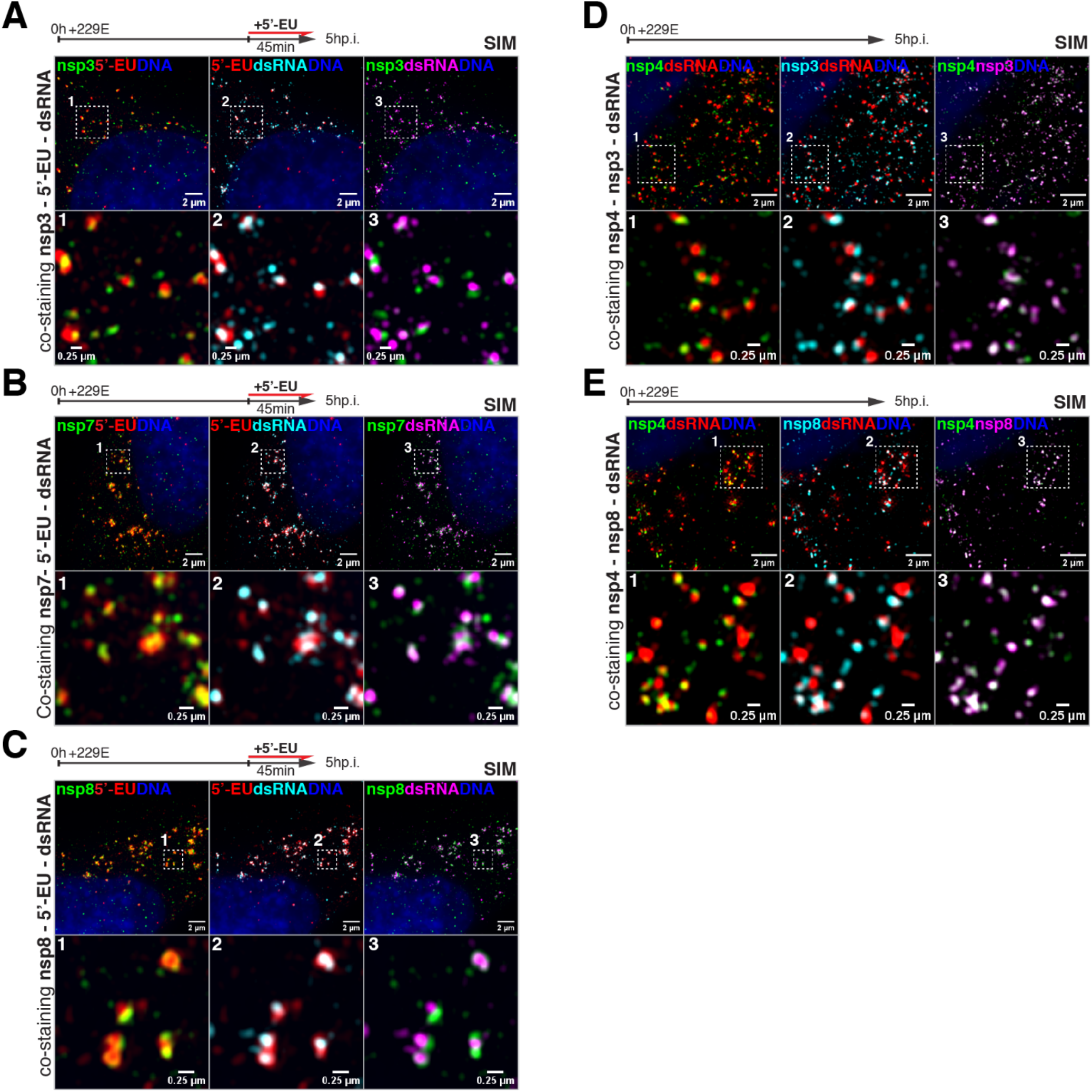
Nsp7 and nsp8 are closely associated with dsRNA and nascent viral RNA. **(A)** hCoV-229E-infected HeLa cells were labeled with EU, then fixed and permeabilized. The cells were reacted by click chemistry with biotin-azide, followed by staining with fluorescent streptavidin. The cells were also co-stained with dsRNA probe and anti-nsp3 antibodies, then imaged by structural illumination microscopy (SIM). **(B)** As in (A), but with staining for nsp7. **(C)** As in (A), but with staining for nsp8. **(D)** As in (A), but with co-staining for nsp4, nsp3 and dsRNA. **(E)** As in (A), but with co-staining for nsp4, nsp8 and dsRNA.

**Fig. S8.**
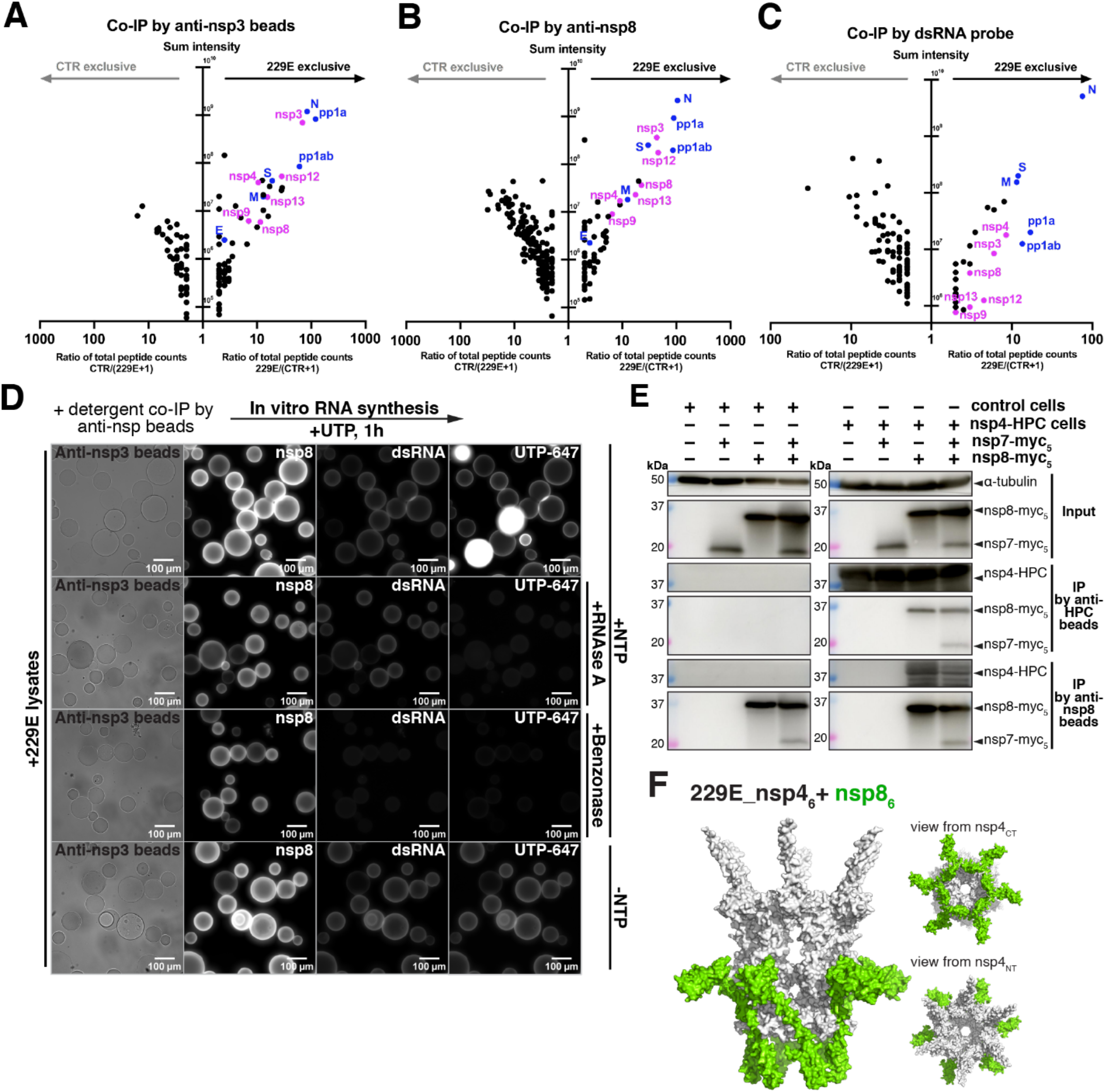
A transcriptionally active replicase-pore complex. **(A-C)** hCoV-229E-infected HeLa cells were lysed, and nuclear pellets were solubilized and subjected to immunoprecipitation with anti-nsp3 antibody beads (A), or anti-nsp8 antibody beads (B), or dsRNA-probe immobilized on beads (C). Co-immunoprecipitated viral proteins were eluted from beads and analyzed by LC-MS/MS. Ratios of total peptide counts in viral-infected samples were calculated by their detected numbers in viral-sample over their detected numbers in control-sample plus one [229E/(CTR+1)]. Ratio of peptide counts for each detected protein (x-axis) was plotted against peptide intensity (y-axis). **(D)** As in (A), but immunoprecipitates on beads were incubated with NTPs and fluorescent UTP; or followed by incubation with enzymes; or with UTP alone. The beads were then fixed and co-stained for dsRNA and nsp8. **(E)** Left: HEK293 cells expressing myc-nsp7, myc-nsp8, or both were lysed and lysates immunoprecipitated with anti-HPC or anti-nsp8 beads. Immunoprecipitated nsps were analyzed by SDS-PAGE and immunoblotting with anti-myc or anti-HPC antibodies. Nsp7 and nsp8 interact. Right: as on the left, but with cells stably expressing nsp4-HPC. Nsp4 co-immunoprecipitates with nsp8 alone, or with co-expressed nsp7-nsp8. **(F)** ColabFold-predicted structure of a complex between hCoV-229E nsp4 hexamer (white) and nsp8 (green). Side view (left) and end views (right) of the molecular surface.

**Fig. S9.**
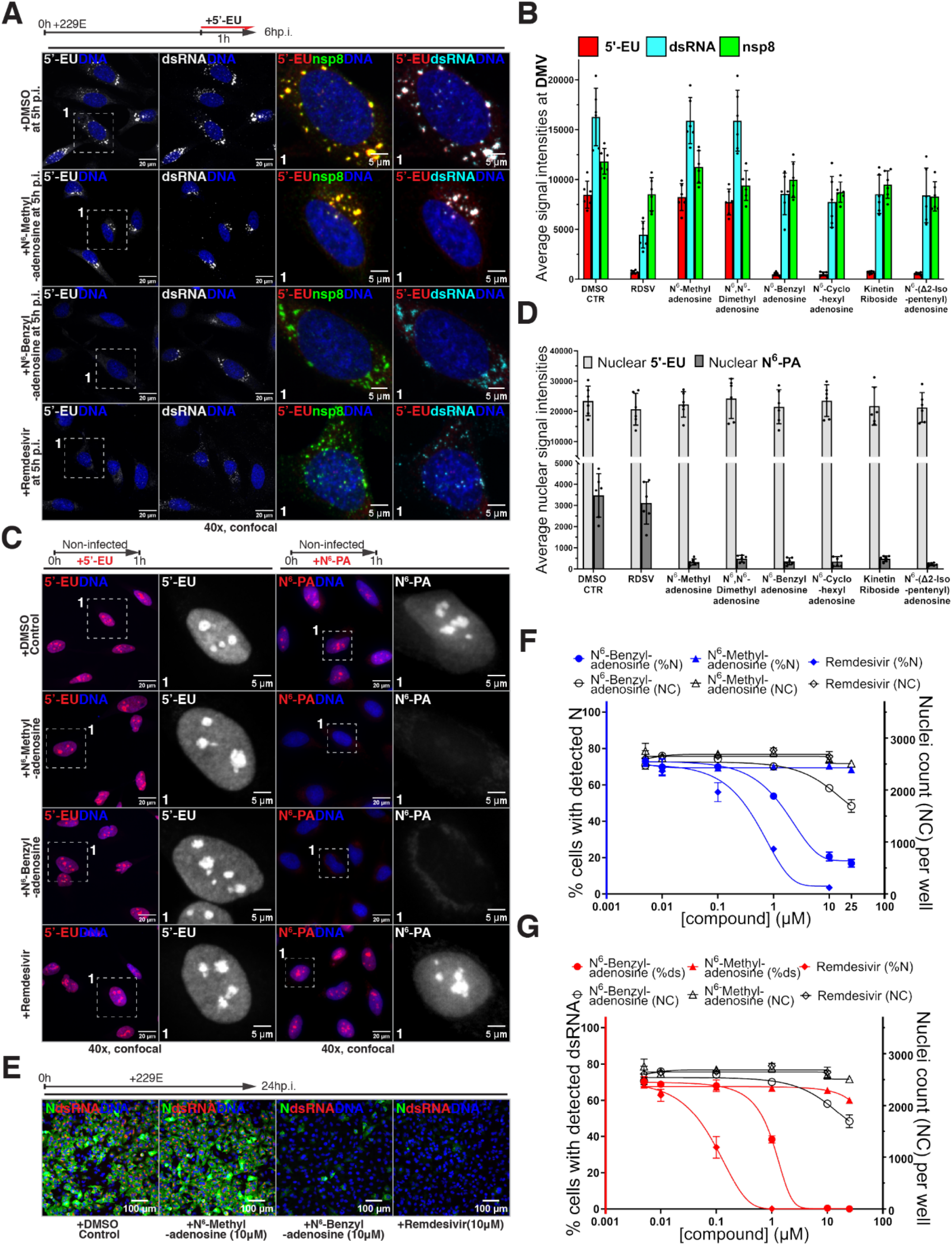
Adenosine analogs with bulky N^6^ modification inhibit viral RNA synthesis and viral replication. **(A)** hCoV-229E-infected HeLa cells were co-incubated with EU and the indicated nucleoside analogs between 5-6h p.i.; and then fixed and processed for detection of EU, dsRNA, and nsp8. **(B)** As in (A), but with additional nucleoside analogs. Average intensities of EU, dsRNA or nsp8 at DMV were quantified. Three non-overlapping fields of view were analyzed per condition, and each condition was repeated twice. (*N*=2, n=6). **(C)** Left: HeLa cells were co-incubated with EU and the indicated nucleoside analogs for 1h, fixed and then stained by click reaction with a fluorescent azide. Right: As on the left, but using N^6^-propargyl-adenosine (N^6^-PA) instead of EU. **(D)** As in (C), but with additional nucleoside analogs. Average intensities of nuclear EU or N^6^-PA were quantified. Three non-overlapping fields of view were analyzed per condition and each condition was repeated 2 times. (*N*=2, n=6). **(E)** hCoV-229E-infected HeLa cells were co-incubated with the indicated compounds for 24hrs, followed by staining with anti-nucleocapsid (N) antibodies and dsRNA probe. **(F and G)** As in (E), but with a dose-response for the indicated compounds. Cells with positive N staining (F), or positive dsRNA staining (G) were quantified at each concentration. N^6^-benzyl-adenosine inhibits viral replication while N^6^-methyl-adenosine does not.

**Fig. S10.**
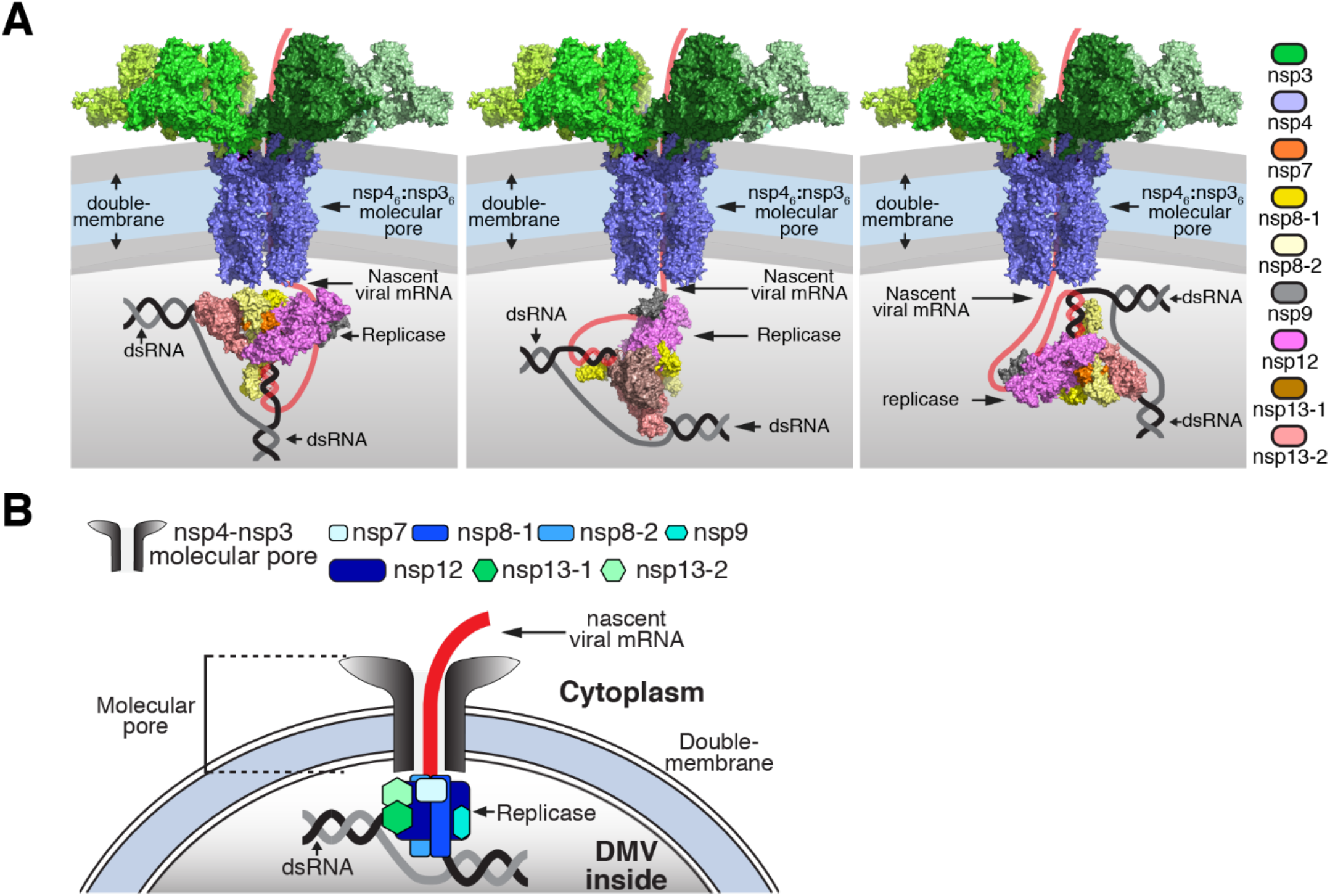
Models of a transcribing replicase-pore complex in the DMV. **(A)** Schematics of a transcribing complex formed by the replicase, pore and dsRNA template. **(B)** Cartoon illustration. The replicase uses dsRNA as a template, and the nascent viral mRNA translocates through the nsp3-nsp4 pore to cytoplasm.

**Table S1.**
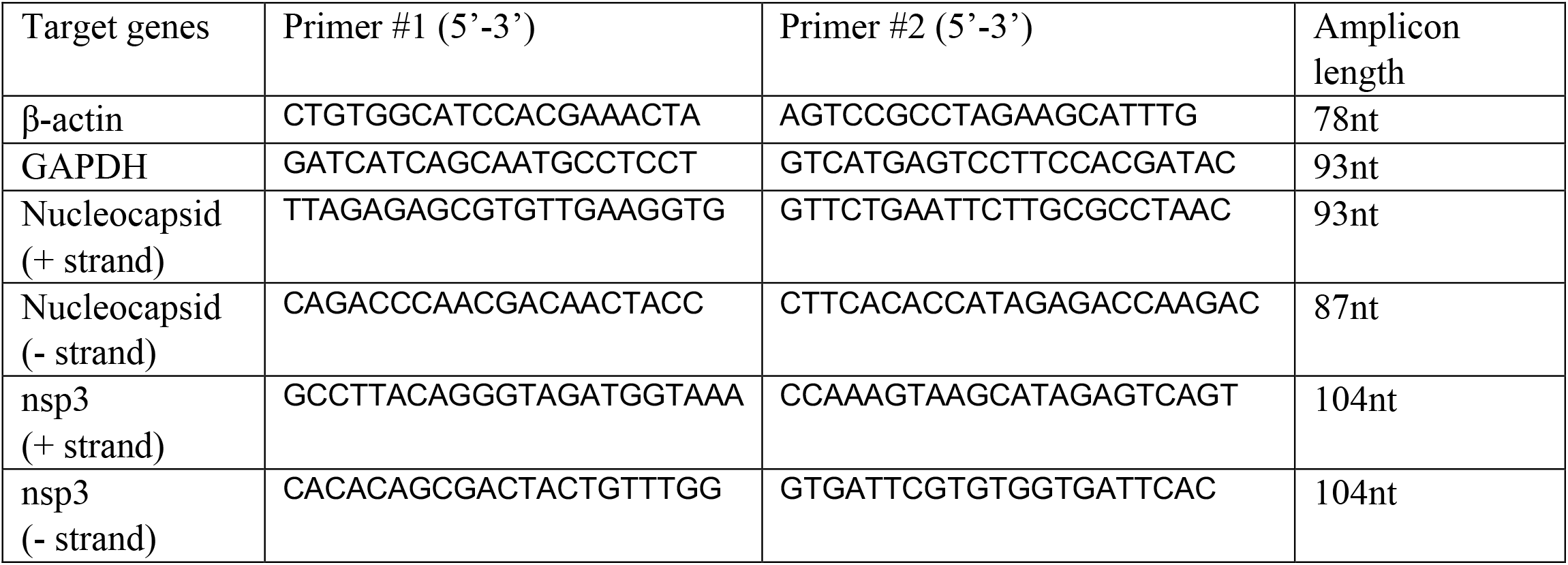
Oligo nucleotide primers used in quantitative PCR in this study.

**Data S1.**
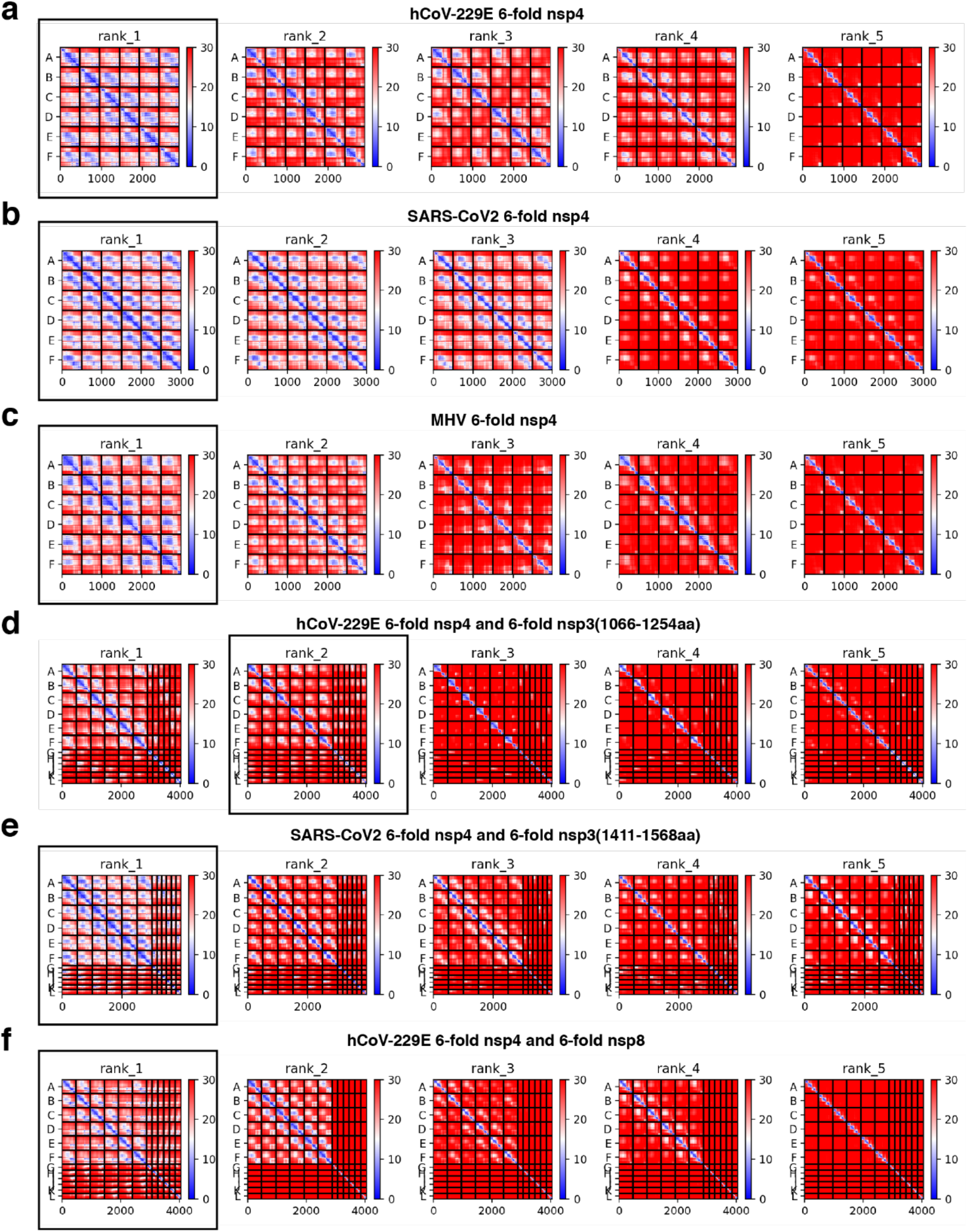
Predicted Alignments Errors (PAE) of structures by ColabFold. **(A)** Predicted Alignments Errors (PAE) of 6-fold hCoV-229E nsp4 by ColabFold. A total of five structures are predicted by ColabFold (from left to right) with their PAE graphs listed from rank 1 to rank 5. A-E indicates six oligomers of nsp4. X axis indicates amino acids positions. The 6-fold nsp4 structure corresponds to rank 1 is used in this study. **(B)** Same as (A), but showing PAE of 6-fold SARS-CoV2 nsp4 structures. The 6-fold nsp4 structure corresponds to rank 1 is used in this study. **(C)** Same as (A), but showing PAE of 6-fold MHV nsp4 structures. The 6-fold nsp4 structure corresponds to rank 1 is used in this study. **(D)** Same as (A), but showing PAE of 6-fold hCoV-229E nsp4 + 6-fold hCoV-229E nsp3(1066-1254aa) structures. The 6-fold nsp4+6-fold nsp3(1066-1254aa) structure corresponds to rank 2 is used in this study. **(E)** Same as (A), but showing PAE of 6-fold SARS-CoV2 nsp4 + 6-fold SARS_CoV2 nsp3(1411-1568aa) structures. The 6-fold nsp4+6-fold nsp3(1411-1568aa) structure corresponds to rank 1 is used in this study. **(F)** Same as (A), but showing PAE of 6-fold hCoV-229E nsp4 + 6-fold hCoV-229E nsp8 structures. The 6-fold nsp4+6-fold nsp8 structure corresponds to rank 1 is used in this study.

